# Functional analysis of the fatty acid and alcohol metabolism of *Pseudomonas putida* using RB-TnSeq

**DOI:** 10.1101/2020.07.04.188060

**Authors:** Mitchell G. Thompson, Matthew R. Incha, Allison N. Pearson, Matthias Schmidt, William A. Sharpless, Christopher B. Eiben, Pablo Cruz-Morales, Jacquelyn M. Blake-Hedges, Yuzhong Liu, Catharine A. Adams, Robert W. Haushalter, Rohith N. Krishna, Patrick Lichtner, Lars M. Blank, Aindrila Mukhopadhyay, Adam M. Deutschbauer, Patrick M. Shih, Jay D. Keasling

## Abstract

With its ability to catabolize a wide variety of carbon sources and a growing engineering toolkit, *Pseudomonas putida* KT2440 is emerging as an important chassis organism for metabolic engineering. Despite advances in our understanding of this organism, many gaps remain in our knowledge of the genetic basis of its metabolic capabilities. These gaps are particularly noticeable in our understanding of both fatty acid and alcohol catabolism, where many paralogs putatively coding for similar enzymes co-exist making biochemical assignment via sequence homology difficult. To rapidly assign function to the enzymes responsible for these metabolisms, we leveraged Random Barcode Transposon Sequencing (RB-TnSeq). Global fitness analyses of transposon libraries grown on 13 fatty acids and 10 alcohols produced strong phenotypes for hundreds of genes. Fitness data from mutant pools grown on varying chain length fatty acids indicated specific enzyme substrate preferences, and enabled us to hypothesize that DUF1302/DUF1329 family proteins potentially function as esterases. From the data we also postulate catabolic routes for the two biogasoline molecules isoprenol and isopentanol, which are catabolized via leucine metabolism after initial oxidation and activation with CoA. Because fatty acids and alcohols may serve as both feedstocks or final products of metabolic engineering efforts, the fitness data presented here will help guide future genomic modifications towards higher titers, rates, and yields.

**IMPORTANCE:** To engineer novel metabolic pathways into *P. putida*, a comprehensive understanding of the genetic basis of its versatile metabolism is essential. Here we provide functional evidence for the putative roles of hundreds of genes involved in the fatty acid and alcohol metabolism of this bacterium. These data provide a framework facilitating precise genetic changes to prevent product degradation and channel the flux of specific pathway intermediates as desired.

## INTRODUCTION

*Pseudomonas putida* KT2440 is an important metabolic engineering chassis, which can readily metabolize compounds derived from lignocellulosic and plastic derived feedstocks (1–3), and has an ever-growing repertoire of advanced tools for genome modification (4–7). Its upper glycolytic pathway architecture enables *P. putida* to natively generate large amounts of reducing equivalent (8), and it more robustly withstands metabolic burdens than many other frequently used host organisms (9). To date, a wide variety of products have been produced through metabolic engineering of *P. putida*, including valerolactam (10), curcuminoids (11), diacids (12), methyl-ketones (13), rhamnolipids (14), cis,cis-muconic acid (15), and many others (16). Recent advances in genome-scale metabolic modeling of *P. putida* make engineering efforts more efficient (7, 17). However, a large gap still exists between genes predicted to encode enzymatic activity and functional data to support these assumptions. Recent characterizations of enzymes and transporters involved in the catabolism of lysine (12, 18), levulinic acid (19), and aromatic compounds (20) highlight the need to continue functionally probing the metabolic capabilities of *P. putida*, because its native catabolism can consume many target molecules and dramatically impact titers.

Amongst the most important metabolisms not yet rigorously interrogated via omics-level analyses are fatty acid and alcohol degradation. Recently, fatty acids have been shown to be a non-trivial component of some feedstock streams (1) and, depending on their chain length, serve as high-value target molecules (21). Furthermore, intermediates in beta-oxidation can be channeled towards mega-synthases to produce more complex molecules (22), or used in reverse beta-oxidation to produce compounds such as medium chain n-alcohols (23). However, assigning the genetic basis of fatty acid degradation is complicated by the presence of multiple homologs of the individual *fad* genes encoded in the genome of *P. putida* KT2440 (17, 24). Although work has been done to either biochemically or genetically demonstrate the substrate specificity of some individual *fad* genes, the majority of these homologs still have no functional data associated with them.

*P. putida* is also able to oxidize and catabolize a wide variety of alcohols. Much work has focused on the unique biochemistry and regulation of two pyrroloquinoline quinone (PQQ)-dependent alcohol dehydrogenases (ADH), *pedE* and *pedH*, which exhibit broad substrate specificity for both alcohols and aldehydes (25, 26). Specific work has also investigated the suitability of *P. putida* for the production of ethanol (27) and the genetic basis for its ability to catabolize butanol and 1,4-butanediol (28–30). *P. putida* is also known for its ability to tolerate solvents and alcohols, making it an attractive host for their industrial production (31, 32). Tolerance to these compounds is a product of both robust efflux pumps (31) and the ability of some strains, such as *P. putida* mt-2, to catabolize a range of organic compounds (33). Metabolic engineering has biologically produced a diverse range of alcohols with a wide array of industrial and commercial uses (34–36). As more alcohol synthesis pathways are engineered into *P. putida*, a more complete understanding of the molecular basis of its catabolic capacities will be required to achieve high-titers.

A recent surge in omics-level data has revealed much about the metabolism of *P. putida*, with adaptive evolution (30), proteomics (10, 28, 29), and ^13^C flux analysis (37–39) yielding valuable insights. An approach that has proven to be particularly powerful is Random Barcode Transposon Sequencing (RB-TnSeq) (40, 41). RB-TnSeq allows rapid and inexpensive genome-wide profiling of individual gene fitness in various conditions, and has been used in *P. putida* to identify numerous novel metabolic pathways and aid in increasing titers of the polymer precursor valerolactam (10, 11, 18–20). Here, we leverage RB-TnSeq to interrogate the genetic basis for the catabolism of multiple fatty acids and alcohols to develop an evidence-based understanding of the enzymes and pathways utilized in these metabolisms.

## RESULTS AND DISCUSSION

### Global Analysis of Fatty Acid Metabolism

To characterize the genetic basis of fatty acid metabolism in *P. putida*, barcoded transposon mutant libraries were grown in minimal media with straight chain fatty acids (C3-C10, C12, and C14), fatty esters (Tween20 and butyl stearate), and an unsaturated fatty acid (oleic acid) as sole carbon sources. Pearson correlation analyses of global fitness patterns revealed that the metabolisms of straight chain fatty acids between C7 and C14 clade together, suggesting similar overall catabolic routes (**Figure 1**). Oleic acid, an 18-carbon monounsaturated fatty acid, also grouped within this clade. Shorter chain fatty acids (<C7) did not show high correlation to one another based on global fitness analyses, suggesting more independent routes of catabolism (**Figure 1**). Annotations in the BioCyc database, functional assignment from a recent metabolic model of *P. putida* KT440 (*i*JN1462), and previous *in vitro* biochemical work predict the existence of several enzymes in the genome of the bacterium that may be putatively involved in fatty acid catabolism: six acyl-CoA ligases, seven acyl-CoA dehydrogenases, seven enoyl-CoA hydratases, four hydroxyacyl-CoA dehydrogenases, and five thiolases (**Figure 2**) (17, 24, 42). Our data show discrete fitness patterns for the steps of beta-oxidation that appear to be largely dictated by chain length (**Figure 2**).

**Figure 1:**
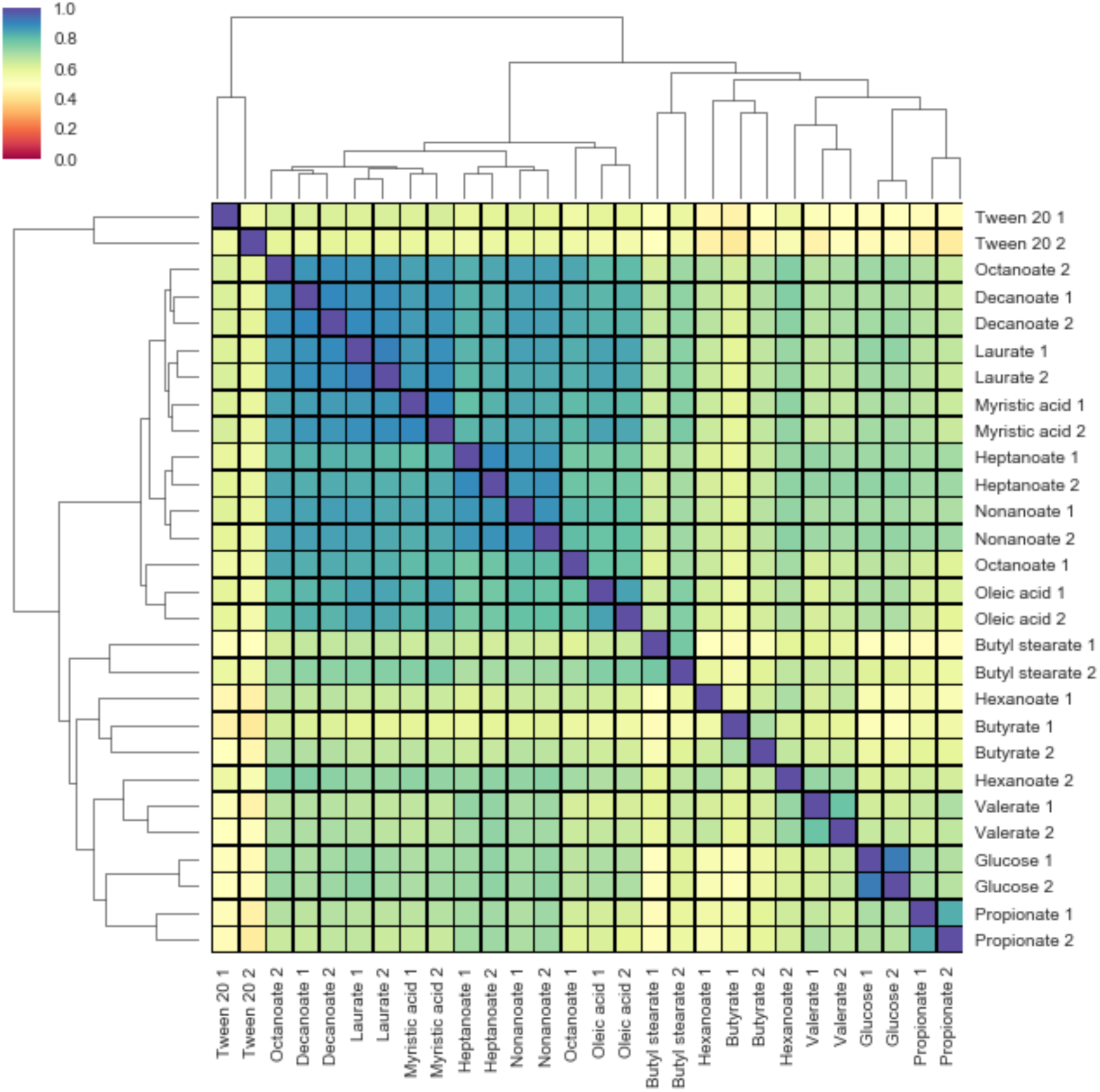
Cladogram correlation matrix of genome-wide fitness data of *P. putida* grown on fatty acids. The matrix shows pairwise comparisons of Pearson correlations of fitness data from *P. putida* KT2440 RB-TnSeq libraries grown on fatty acids as well as glucose. The legend in top left shows Pearson correlation between two conditions with blue showing *r* = 1, and red showing *r =* 0. The conditions were tested in duplicate and the data from each are numbered (1 & 2).

**Figure 2:**
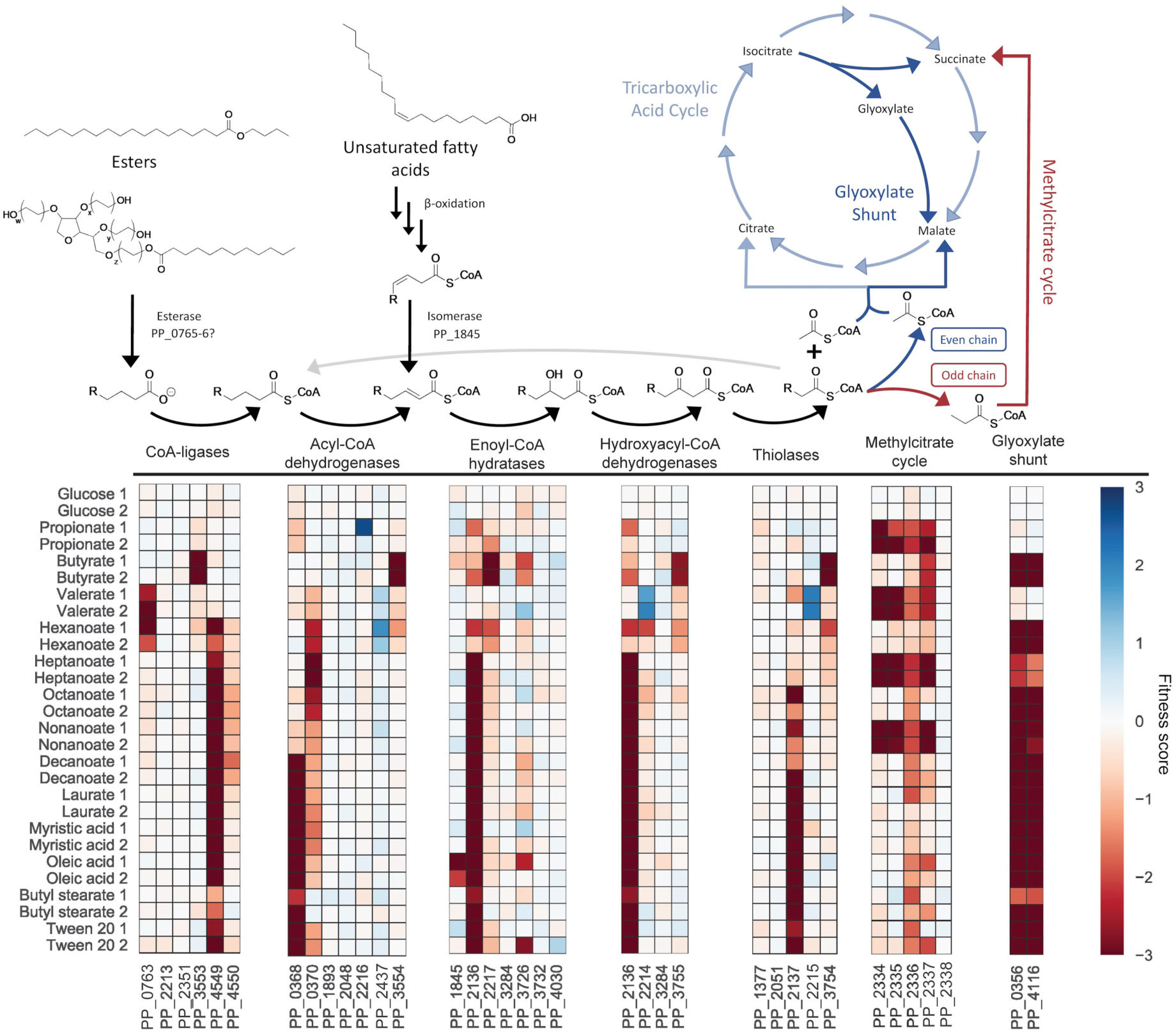
Overview of fatty acid catabolic pathways of *P. putida* KT2440. The above diagram shows the catabolic steps of fatty ester and saturated/unsaturated fatty acid catabolism in *P. putida* KT2440, in addition to their connections to the glyoxylate shunt and the methylcitrate cycle. The heatmaps below show fitness scores when grown on fatty acids or glucose for the specific genes proposed to catalyze individual chemical reactions. Colors represent fitness scores, with blue representing positive fitness and red representing negative fitness.

When grown on fatty acids, many bacteria require the anaplerotic glyoxylate shunt to avoid depleting TCA cycle intermediates during essential biosynthetic processes. In *P. putida*, the two steps of the glyoxylate shunt are encoded by PP_4116 (*aceA* - isocitrate lyase) and PP_0356 (*glcB* - malate synthase). Transposon mutants in both of these genes showed serious fitness defects (fitness score < −3) when grown on nearly all of the fatty acids tested (**Figure 2**). However, the glyoxylate shunt genes appeared dispensable for growth on valerate (C5), and showed a more severe fitness defect when grown on heptanoate (C7). Complete beta-oxidation of valerate and heptanoate results in ratios of propionyl-CoA to acetyl-CoA of 1:1 and 1:2, respectively. This higher ratio of 3-carbon to 2-carbon production presumably offers an alternate means to replenish TCA cycle intermediates in the absence of a glyoxylate shunt (**Figure 2**).

In order to utilize the propionyl-CoA generated by beta-oxidation of odd-chain fatty acids, bacteria often employ the methylcitrate cycle (MCC), producing succinate and pyruvate from oxaloacetate and propionyl-CoA. In *P. putida*, the MCC is catalyzed via methylcitrate synthase (*prpC* - PP_2335), 2-methylcitrate dehydratase (*prpD* - PP_2338, or *acnB* - PP_2339), aconitate hydratase (*acnB* - PP_2339, or *acnA2* - PP_2336), and 2-methylisocitrate lyase (*mmgF* - PP_2334) (**Supplementary Figure 1**). Unsurprisingly, the MCC appeared to be absolutely required for growth on propionate (C3), valerate (C5), heptanoate (C7), and nonanoate (C9), with PP_2334, PP_2335, and PP_2337 (a putative AcnD-accessory protein) showing severe fitness defects (**Figure 2, Supplementary Figure 1**). While PP_2338 (*prpD*) encodes for a 2-methylcitrate dehydratase, transposon mutants showed no fitness defects when grown on odd-chain fatty acids. This reaction is likely carried out by PP_2339 (*acnB* - a bifunctional 2-methylcitrate dehydratase/aconitase hydratase B); however, there were no mapped transposon insertions for this gene (**Figure 2, Supplementary Figure 1**). This suggests that PP_2339 was essential during the construction of the RB-TnSeq library. Furthermore, PP_2336 showed relatively modest fitness defects when grown on propionate and other odd-chain fatty acids, suggesting that PP_2339 likely accounts for much of the methylaconitate hydratase activity as well (**Figure 2, Supplementary Figure 1**).

### Long and Medium Chain Fatty Acid Catabolism

Pearson correlation analysis of fitness data indicated that both long and medium chain fatty acids are likely catabolized via similar pathways. Fitness data suggests that FadD1 (PP_4549) catalyzes the initial CoA-ligation of C7 to C18 fatty acids, and may potentially act on C6 as well (**Figure 2**). Disruption of *fadD2* (PP_4550) did not cause fitness defects as severe as those seen in *fadD1* mutants, although it did result in moderate fitness defects when grown on C8-C10 fatty acids. These data are consistent with the biochemical characterization of FadD1 from *P. putida* CA-3, which showed greater activity on longer chain alkanoic and phenylalkanoic acids than on shorter chain substrates (43). For fatty acids with chain lengths of C10 and greater, the data suggest that the *fadE* homolog PP_0368 is the primary acyl-coA dehydrogenase, while the nearby *fadE* homolog PP_0370 appears to be preferred for C6-C8 fatty acids (**Figure 2**). A relatively even fitness defect for these two *fadE* homologs indicates that PP_0368 and PP_0370 may have equal activity on nonanoate (**Figure 2**). These data are supported by a previous biochemical characterization of PP_0368, in which it showed greater activity on chain lengths longer than C9 (44). The *fadB* homolog PP_2136 showed severe fitness defects when grown on all fatty acids with chain lengths of C6 and longer, implicating it as the primary enoyl-CoA hydratase/3-hydroxy-CoA dehydrogenase for those substrates (**Figure 2**). *P. putida* was able to grow on the unsaturated substrate oleic acid, and is likely able to isomerize the position of the unsaturated bond via the enoyl-CoA isomerase PP_1845, which showed specific fitness defects when grown on oleic acid (**Figure 2**). *P. putida*’s primary long chain thiolase appears to be the *fadA* homolog PP_2137, which showed severe to moderate fitness defects when grown on fatty acids with chain lengths C8 or longer (**Figure 2**). Fitness data for mutant pools grown on heptanoate showed minor fitness defects for both PP_2137 and PP_3754 (*bktB*), suggesting that both thiolases may work on C7 substrates (**Figure 2**).

Both long chain fatty esters tested (Tween 20 and butyl stearate) appeared to utilize the same *fad* homologs as the long chain fatty acids. However, before either molecule can be directed towards beta-oxidation, Tween 20 and butyl stearate must be hydrolyzed to generate a C12 or C18 fatty acid, respectively. To date, no such hydrolase has been identified in *P. putida* KT2440. Comparing the mutant fitness scores between Tween 20 and laurate (C12) carbon source experiments revealed six genes (PP_0765, PP_0766, PP_0767, PP_0914, PP_2018, and PP_2019) that had significant and severe fitness defects specific to Tween 20 (fitness score < −2, *t* > |4|) in both biological replicates (**Figure S2**). The same comparison between butyl stearate and myristate (C14) revealed four genes specific to the fatty ester (PP_0765, PP_0766, PP_2018, and PP_4058) that had significant severe fitness defects (fitness score < −2, *t* > |4|) in both biological replicates (**Figure S2**). As PP_0765-6 and PP_2018 appear to have specific importance in both of the ester conditions tested, it may be possible that they contribute to the hydrolysis of the fatty ester bonds. However, it is also possible that the esterase is secreted or associated with the outer membrane (45), in which case its enzymatic activity would be shared amongst the library and it would not have the associated fitness defect expected (10).

The genes PP_2018 and PP_2019 encode a BNR-domain containing protein and a RND-family efflux transporter, respectively, and are likely co-expressed in an operon that also includes PP_2020 and PP_2021. Interestingly, although PP_2021 codes for a putative lactonase, transposon mutants had no apparent fitness defect with either of the fatty esters as the carbon source. PP_0765 and PP_0766 encode a DUF1302 family protein and DUF1329 family protein, respectively. Given their similar fitness scores, they are likely coexpressed in an operon positively regulated by the LuxR-type regulator PP_0767 (**Figure S2**). Previous work in multiple other species of *Pseudomonas* has observed cofitness of DUF1302/DUF1329 family genes with BNR-domain and RND-family efflux genes when grown on Tween 20 (41). The authors proposed that these genes may work together in order to export a component of the cell wall. However, an alternative hypothesis could be that PP_0765 and PP_0766 contribute to catalyzing the hydrolysis of fatty esters, accounting for the missing catabolic step of butyl stearate and Tween 20. This hypothesis is bolstered somewhat by the co-localization of PP_0765/PP_0766 near fatty acid catabolic genes in *P. putida* KT2440 and many other *Pseudomonads* (**Figure S3**). Future work will need to be done to biochemically characterize the potential enzymatic activity of these proteins.

### Short Chain Fatty Acid Catabolism

In our genome-wide fitness assays, the mutant fitness patterns of C6 or shorter fatty acid carbon sources had lower Pearson correlation between one another than the correlations within long and medium-chain fatty acids (**Figure 1**). These global differences reflect what appear to be discrete preferences in beta-oxidation enzymes for growth on short chain fatty acids. Fitness data suggest that while both CoA-ligases PP_0763 and PP_4559 are required for growth on hexanoate, only PP_0763 is required for growth on valerate (**Figure 2**). Furthermore, the putative positive regulator of PP_0763, LuxR-family transcription factor PP_0767, also showed a significant fitness defect (−2.0) when grown on both valerate and hexanoate (**Figure 2**). PP_0370 seems to be the acyl-CoA dehydrogenase largely responsible for hexanoate catabolism, though PP_3554 mutants also have minor fitness defects. The dehydrogenation of valeryl-coA appears to be distributed between the activities of PP_0368, PP_0370, and PP_3554, with no single acyl-CoA dehydrogenase mutant demonstrating a strong fitness defect when grown on valerate (**Figure 2**). Interestingly, though previous biochemical analysis had demonstrated that PP_2216 has activity on C4-C8 acyl-CoA substrates with a preference for shorter chain lengths (46), we observed no fitness defects for PP_2216 mutants when grown on any fatty acid carbon source (**Figure 2**).

It appears that the role of enoyl-CoA hydratase or hydroxyacyl-CoA dehydrogenase may be distributed across multiple enzymes for both hexanoate and valerate. Growth on hexanoate resulted in moderate fitness defects in mutants disrupted in the predicted enoyl-CoA hydratases PP_2136, PP_2217, and PP_3726; however, for mutants grown on valerate, there were almost no observable fitness defects for any of the enoyl-CoA hydratase enzymes examined in the study, suggesting that for this chain length significant functional complementation exists between the *fadB* homologs (**Figure 2**). Fitness data suggest that PP_2136 (*fadB*), PP_2214 (a predicted type II 3-hydroxyacyl-CoA dehydrogenase), and PP_3755 (a 3-hydroxybutyryl-CoA dehydrogenase) may all be involved in the dehydrogenation of 3-hydroxyhexanoyl-CoA (**Figure 2**), while there appears to be a distribution of *fadB*-like activity when it comes to the dehydrogenation of 3-hydroxyvaleryl-CoA, with PP_3755 showing only a slight fitness defect on valerate. Intriguingly, mutants disrupted in the predicted type-2 acyl-CoA dehydrogenase PP_2214 showed apparent increased fitness when grown on valerate (**Figure 2**). As with heptanoate, fitness data from mutant pools grown on valerate or hexanoate suggest that both PP_2137 and PP_3754 may catalyze the terminal thiolase activity of these substrates. The lack of pronounced fitness phenotypes for the beta-oxidation steps of both valerate and hexanoate underscores the necessity for further *in vitro* biochemical interrogation of these pathways.

Both the butanol and butyrate metabolism of *P. putida* have been studied in detail through omics-level interrogation across multiple strains (28, 29). Previous work showed that during growth on n-butanol, which is later oxidized to butyrate, three CoA-ligases are up-regulated: PP_0763, PP_3553, and PP_4487 (*acsA-1* - an acyl-CoA synthase) (29). However, our butyrate carbon source experiments only revealed strong fitness defects in PP_3553 mutants (**Figure 2**, **Figure 3A**). The same work found that PP_3554 was the only upregulated acyl-CoA dehydrogenase, which agrees with the strong fitness defect we observed in mutants of that gene (29). That prior work did not find upregulation of any enoyl-CoA hydratase in *P. putida* grown on butanol, but this is likely reflective of redundancy in this step; we observed fitness defects in multiple genes, including PP_2136, PP_2217, and PP_3726, with mutants in PP_2217 demonstrating the most severe fitness defect (**Figure 2, Figure 3A**). Hydroxyacyl-CoA dehydrogenase PP_2136 and 3-hydroxybutyryl-CoA dehydrogenase PP_3755 (*hbd*) have both been shown to be upregulated during growth on butanol (29). While our data showed fitness defects in both of these genes, the defect of PP_3755 mutants was much more severe. Three different thiolases (PP_2215, PP_3754, and PP_4636) and the 3-oxoacid CoA-transferase *atoAB* were previously observed to be upregulated during growth on butanol, but of these genes, only PP_3754 (*bktB*) had a strong fitness defect, implying that it is the main thiolase for the terminal step of butyrate catabolism (**Figure 2, Figure 3A**).

**Figure 3:**
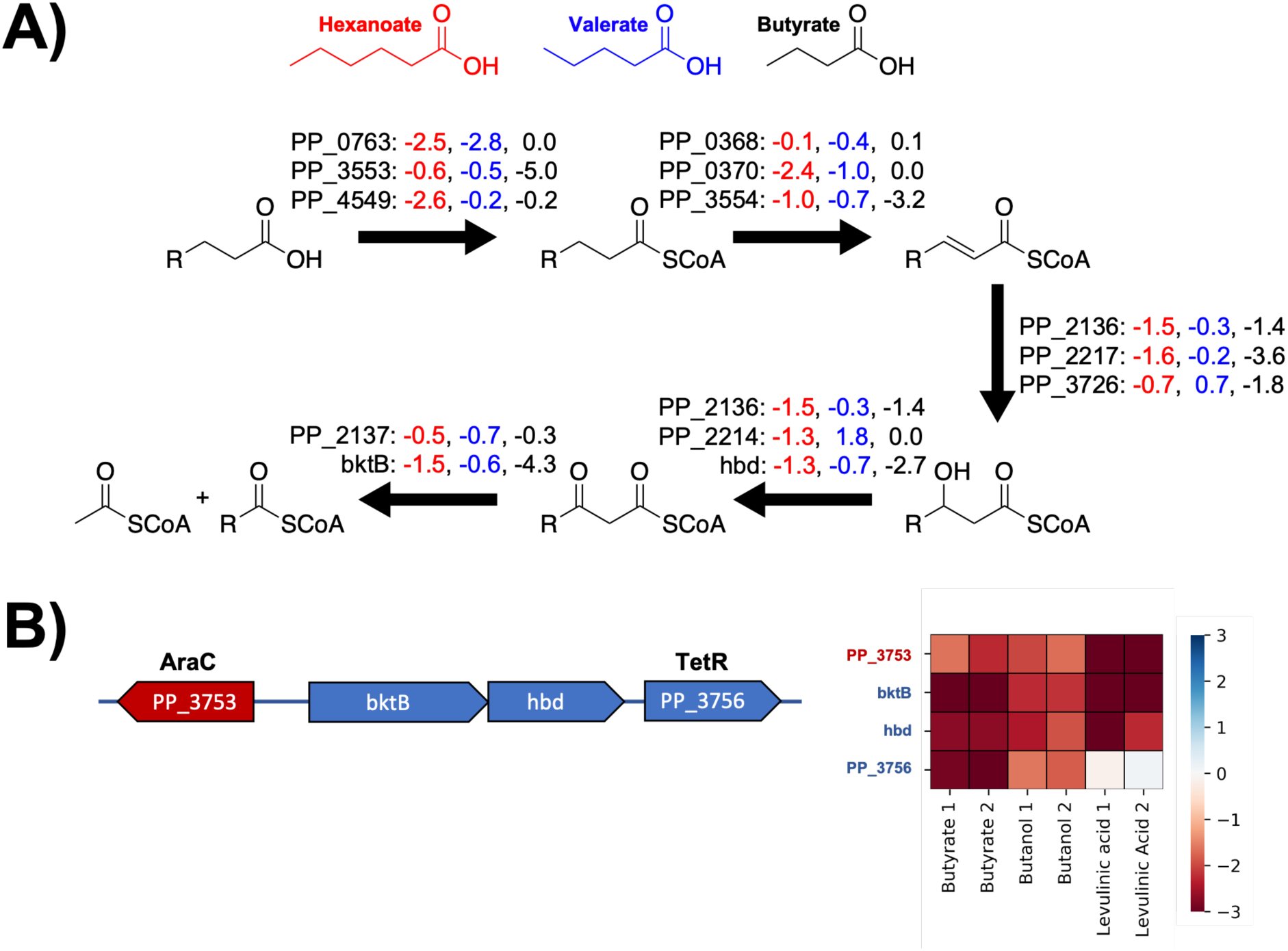
Putative pathways for short chain fatty acid catabolism in *P. putida* KT2440. A) Individual enzymatic steps that potentially catalyze the steps of beta-oxidation for short chain fatty acids, fitness scores are listed to the right of each enzyme when grown on either butyrate, valerate, or hexanoate. B) The operonic structure of *btkB* and *hdb* flanked by an AraC-family (PP_3753) and TetR-family (PP_3756). The heatmap shows fitness scores of the genes when grown on butyrate, butanol, or levulinic acid.

The inability of the RB-TnSeq data to clearly show which enzymes are likely responsible for specific beta-oxidation reactions suggest multiple enzymes may catalyze these steps. In addition to the lack of genotype to phenotype clarity in the enzymes responsible for the catabolic steps, we observed additional phenotypes within our fitness data that portray a complex picture of short chain fatty acid metabolism in *P. putida*. The TetR-family repressor *paaX* (PP_3286) was shown to have a negative fitness score when mutant pools were grown on fatty acids with chain lengths C7 or below (**Figure S4**). PaaX negatively regulates the *paa* gene cluster encoding the catabolic pathway for phenylalanine (47, 48), implying that presence of phenylalanine catabolism impedes growth on short chain fatty acids. It is therefore somewhat surprising that no individual mutant within the *paa* gene cluster shows a fitness increase when grown on short chain fatty acids, though no robust fitness data exists for *paaJ* (PP_3275 - a 3-oxo-5,6-didehydrosuberyl-coA thiolase) (**Figure S4**).

Mutants in MerR-family regulator PP_3539 showed very high fitness benefits (fitness scores of 3.8 and 4.7 in two biological replicates) when grown on valerate. PP_3539 likely increases expression of *mvaB* (PP_3540 - hydroxymethyl-glutaryl-CoA lyase), thus suggesting that decreased levels of MvaB activity may benefit *P. putida* valerate catabolism. Unfortunately, there are no fitness data available for *mvaB*, likely because it is essential under the conditions in which the initial transposon library was constructed. The genes *hdb* and *bktB*, encoding the terminal two steps of butyrate metabolism, are flanked upstream by an AraC-family regulator (PP_3753) and downstream by a TetR-family regulator (PP_3756); the latter is likely co-transcribed with the butyrate catabolic genes (**Figure 3B**). When grown on butyrate, mutants in both PP_3753 and PP_3756 show decreased fitness; however, previous work to evaluate global fitness of *P. putida* grown on levulinic acid showed negative fitness values only for PP_3753, *htb*, and *btkB* (**Figure 3B**). These results suggest that the TetR repressor may be responding to a butyrate specific metabolite. Finally, across multiple fitness experiments, the TonB siderophore receptor PP_4994 and the TolQ siderophore transporter PP_1898 showed fitness advantages when grown on fatty acids, especially on hexanoate (**Figure S5**). Together, these results suggest that a wide range of environmental signals impact how *P. putida* is able to metabolize short chain fatty acids.

### Global Analysis of Alcohol Catabolism

In addition to its ability to robustly catabolize a wide range of fatty acid substrates, *P. putida* is also capable of oxidizing and catabolizing a wide variety of alcohols into central metabolism through distinct pathways. To further our understanding of these pathways, transposon libraries were grown on a number of short n-alcohols (ethanol, butanol, and pentanol), diols (1,2-propanediol, 1,3-butanediol, 1,4-butanediol, and 1,5-pentanediol), and branched chain alcohols (isopentanol, isoprenol, and 2-methyl-1-butanol). Relative to growth on fatty acids, fitness experiments of *P. putida* grown on various alcohols showed less correlation to one another, reflecting the more diverse metabolic pathways used for their catabolism (**Figure 4A**). The initial step of the catabolism of many primary alcohols is the oxidation of the alcohol to its corresponding carboxylic acid. The BioCyc database features 14 genes annotated as alcohol dehydrogenases (PP_1720, PP_1816, PP_2049, PP_2492, PP_2674, PP_2679, PP_2682, PP_2827, PP_2953, PP_2962, PP_2988, PP_3839, PP_4760, and PP_5210) (24). Fitness data showed that the majority of these alcohol dehydrogenases had no fitness defects when grown on the alcohols used in this study (**Figure 4B**).

**Figure 4:**
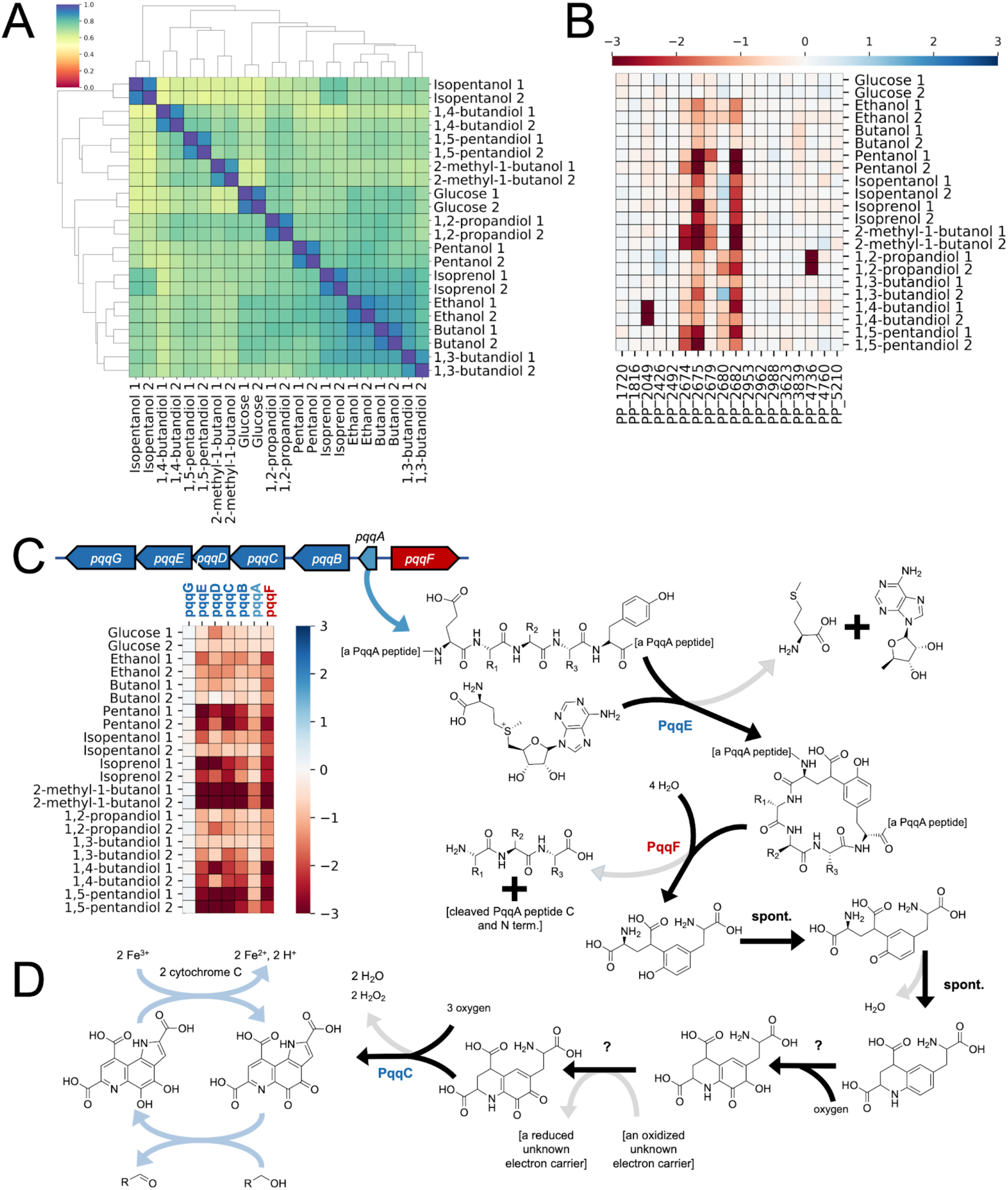
Global analysis of alcohol metabolism in *P. putida*. A) Pairwise comparisons of Pearson correlations of fitness data from *P. putida* KT2440 RB-TnSeq libraries grown on alcohols as well as glucose grouped by overall similarity. Colors bar at top left shows the Pearson coefficient with 1 indicating greater similarity and 0 indicating greater dissimilarity. B) Heatmap shows the fitness scores of all alcohol dehydrogenases annotated on the BioCyc database as well as the cytochrome C PP_2675 when grown on various alcohols and glucose. C) Operonic diagram of the *pqq* cluster in *P. putida* and the corresponding biosynthetic pathway for the PQQ cofactor and D) How PQQ cofactors are regenerated by cytochrome C. Heatmap shows fitness scores for individual *pqq* cluster genes when grown on alcohols and glucose.

The alcohol dehydrogenases that showed the most consistent fitness defects across multiple conditions were the two PQQ-dependent alcohol dehydrogenases PP_2674 (*pedE*) and PP_2679 (*pedH*), as well as the Fe-dependent alcohol dehydrogenase PP_2682 (*yiaY*) (**Figure 4B**). Both *pedE* and *pedH* have been extensively studied in *P. putida* and other related bacteria, and are known to have broad substrate specificities for alcohols and aldehydes (25, 26, 49). Their activity is dependent on the activity of *pedF* (PP_2675), a cytochrome *c* oxidase that regenerates the PQQ cofactor (25). In *P. aeruginosa*, a homolog of *yiaY* (*ercA*) was shown to have a regulatory role in the expression of the *ped* cluster, and was not believed to play a direct catabolic role (50). In most conditions tested, disruption of *pedF* caused more severe fitness defects than either *pedE* or *pedH* individually, suggesting they can functionally complement one another in many cases. However, growth on 2-methyl-1-butanol and 1,5-pentanediol both showed more severe fitness defects in *pedE* mutants compared to *pedF* (**Figure 4B**). In many other alcohols, including ethanol and butanol, even disruption of *pedF* did not cause extreme fitness defects, suggesting the presence of other dehydrogenases able to catalyze the oxidation (**Figure 4B**).

The transcriptional regulatory systems that activate expression of various genes in the *ped* cluster could also be identified from these data. Mutants in either member of the sensory histidine kinase/response regulator (HK/RR) two component system, *pedS2*/*pedR2*, showed significant fitness defects when 2-methyl-1-butanol was supplied as the sole carbon source. This HK/RR signaling system has been shown to activate the transcription of *pedE* and repress *pedH* in the absence of lanthanide ions (51). Since lanthanides were not supplied in the medium, this likely explains the fitness defect observed in *pedS2/pedR2*. The transcription factor *pedR1* (*agmR*) was also found to affect host fitness when grown on various alcohols (**Figure 5**). This gene has been identified in *P. putida* U as an activator of long chain (C6+) n-alcohol and phenylethanol catabolism (52). In *P. putida* KT2440, *pedR1* has been associated with the host response to chloramphenicol, and its regulon has been elucidated previously (53). Our data reflect the literature, indicating that *pedR1* functions as a transcriptional activator of the *ped* cluster and *pedR2* functions as a specific regulator of *pedE* and *pedH*.

**Figure 5:**
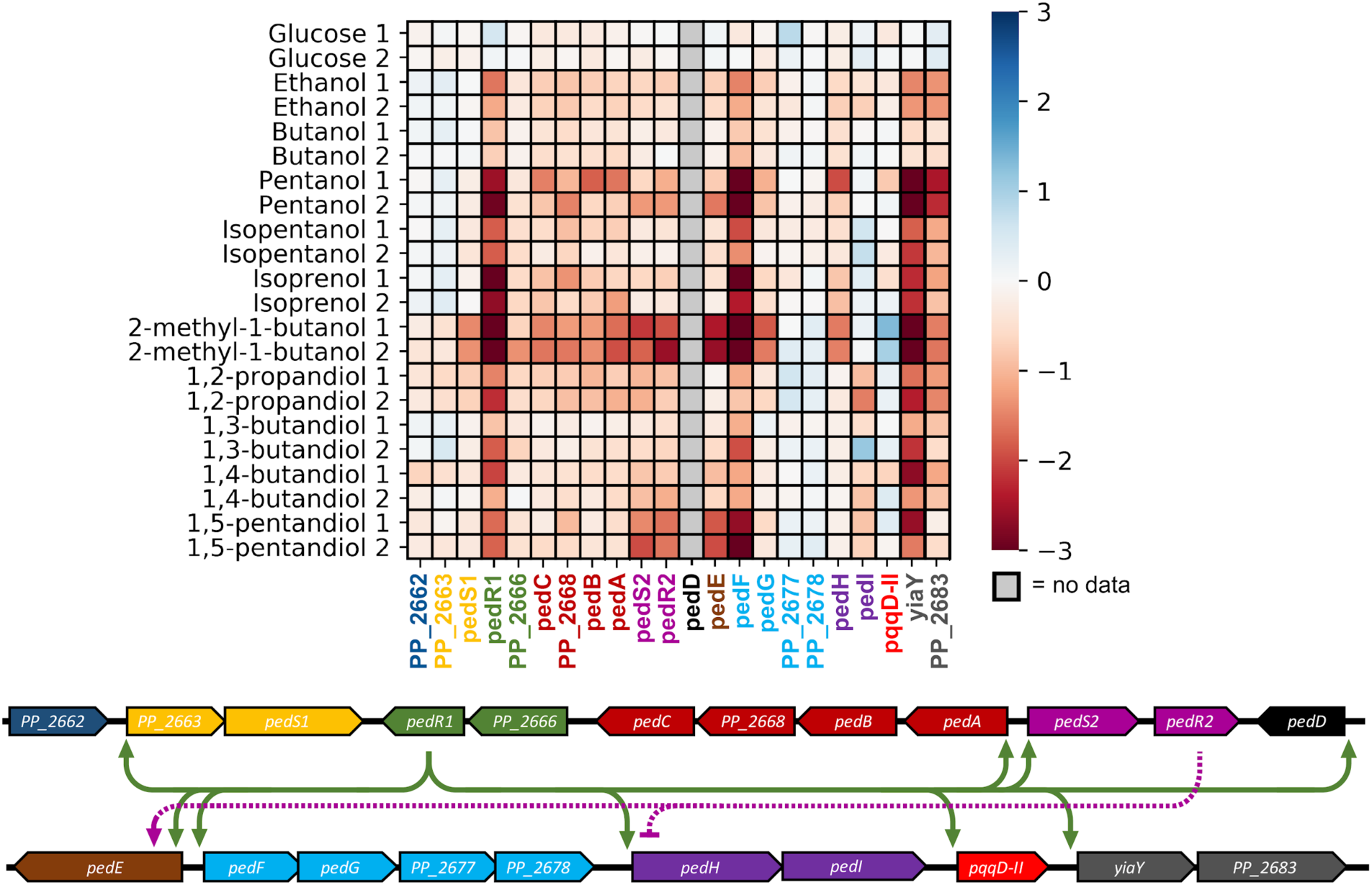
Essentiality and regulation of the *ped* cluster. (Top) Heatmap depicting the fitness scores for genes in the *ped* cluster (PP_2662 to PP_2683) during growth on various short chain alcohols. (Bottom) Genomic context for the *ped* cluster in *P. putida* KT2440. Arrows depict transcriptionally upregulated genes of *pedR1* and *pedR2*. Blunt arrows point to genes predicted to be transcriptionally repressed in the condition tested.

Unsurprisingly, the genes required for the biosynthesis of the PQQ cofactor were also amongst the most co-fit (cofitness is defined as Pearson correlation between fitness scores of two genes over many independent experimental conditions) with both *pedF* and *yiaY*. *P. putida* synthesizes PQQ via a well-characterized pathway, starting with a peptide encoded by the gene *pqqA* (PP_0380) which is then processed by *pqqE*, *pqqF*, and *pqqC* to generate the final cofactor (**Figure 4C**). The three synthetic genes (*pqqEFC*) all showed significant fitness defects on the same alcohols as the *pedF* mutants, while *pqqA* showed a less severe fitness phenotype (**Figure 4C**). However, the small size of *pqqA* resulted in few transposon insertions, making it difficult to draw confident conclusions. Two genes showed similar defective fitness patterns on select alcohols: *pqqB*, which has been proposed to be an oxidoreductase involved in PQQ biosynthesis; and *pqqD*, a putative PQQ carrier protein. Previous work regarding a PqqG homolog from *Methylorubrum extorquens* suggested that it forms a heterodimeric complex with PqqF that proteolytically processes PqqA peptides, although PqqF was sufficient to degrade PqqA on its own (54). Fitness data from *P. putida* may support this hypothesis, as there was no observed fitness defect in *pqqG* mutants when grown on any alcohol, suggesting that the bacterium is still able to process PqqA with PqqF alone (**Figure 4C**).

### Short chain alcohol metabolism

The metabolism of n-alcohols almost certainly proceeds through beta-oxidation using the same enzymatic complement as their fatty acid counterparts. This relationship is reflected in the high correlation in global fitness data between alcohols and fatty acids of the same chain length (ethanol and acetate - *r* = 0.72, butanol and butyrate - *r* = 0.66, pentanol and valerate - *r* = 0.72). However, given previous work and our fitness data, the initial oxidation of these alcohols appears to be quite complex. Biochemical characterization of both PedE and PedH have shown that both have activity on ethanol, acetaldehyde, butanol, butyraldehyde, hexanol, and hexaldehyde (25). When grown on n-pentanol, mutants disrupted in *pedF* show severe fitness defects, suggesting that PedH and PedE are the primary dehydrogenases responsible for pentanol oxidation (**Figure 4B**, **Figure 5A**). However, when grown on either ethanol or n-butanol, both the PQQ-dependent alcohol dehydrogenases (PQQ-ADHs) and *pedF* show less severe fitness defects compared to when they are grown on pentanol (**Figure 4B**). This implies that other dehydrogenases are also capable of these oxidations. One likely candidate may be PP_3839, which shows a minor fitness defect when grown on n-butanol and has been biochemically shown to oxidize coniferyl alcohol (**Figure 4B**) (55). Individual gene deletion mutants of either *pedF* (PP_2675) or PP_3839 showed only minor growth defects when grown on either ethanol, butanol, or pentanol as a sole carbon source (**Figure 7**). However, when both genes were deleted, no growth was observed on these substrates, suggesting that the PQQ-ADHs and PP_3839 are the primary dehydrogenases responsible for the oxidation of short chain n-alcohols (**Figure 7**).

**Figure 6:**
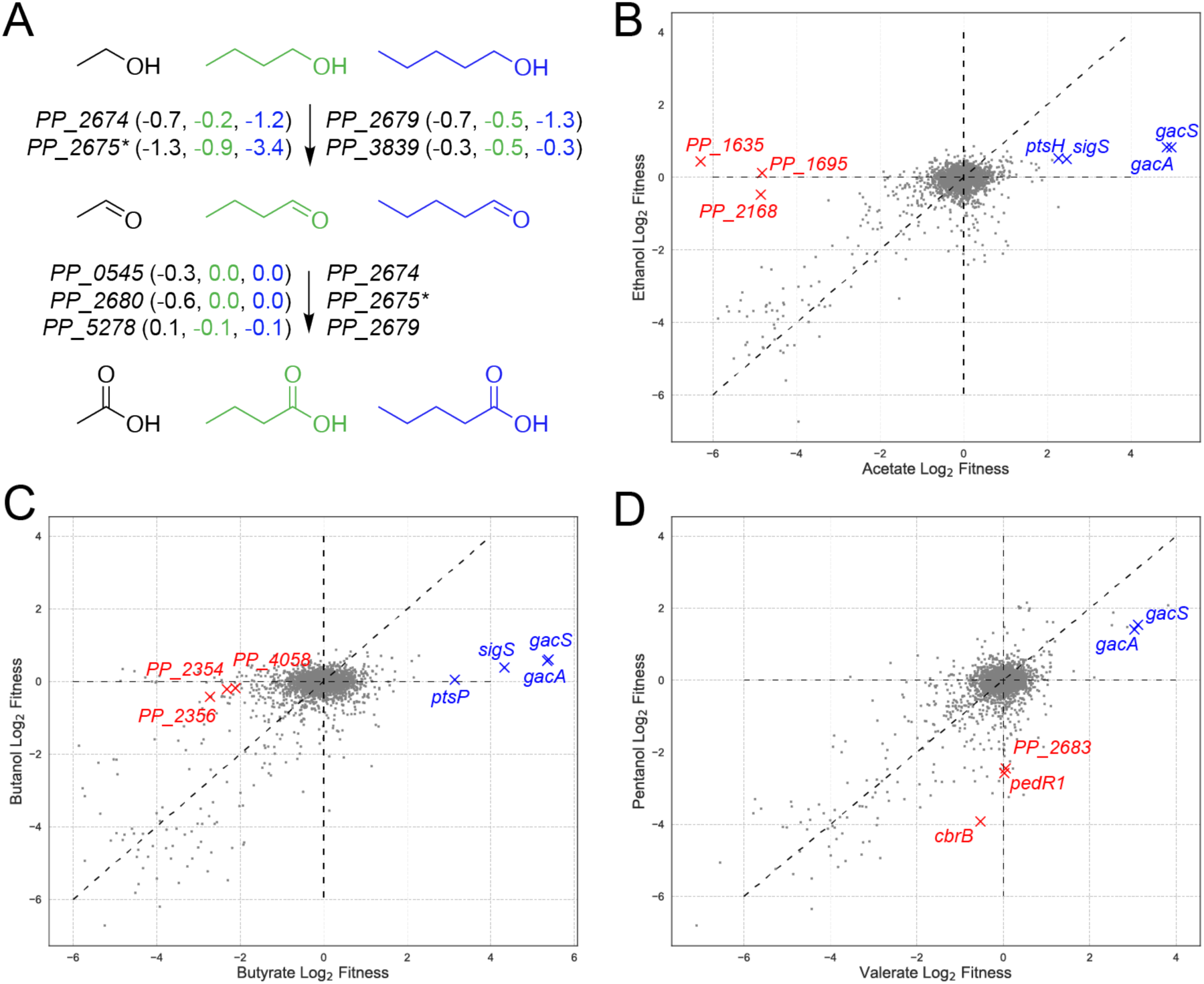
Analysis of short chain alcohol metabolism in *P. putida*: A) Putative genes involved in the initial oxidation steps of short chain alcohol assimilation in *P. putida*. PP_2675 (PedF) is involved in the regeneration of the PQQ cofactor predicted to be necessary for these oxidation reactions of PP_2764 (PedE) and PP_2769 (PedH). Average fitness scores for two biological reps are shown next to each gene for ethanol (black), butanol (green), and pentanol (blue). Scatter plots show global fitness scores for ethanol versus acetate (B), butanol versus butyrate (C), and pentanol versus valerate (D).

**Figure 7:**
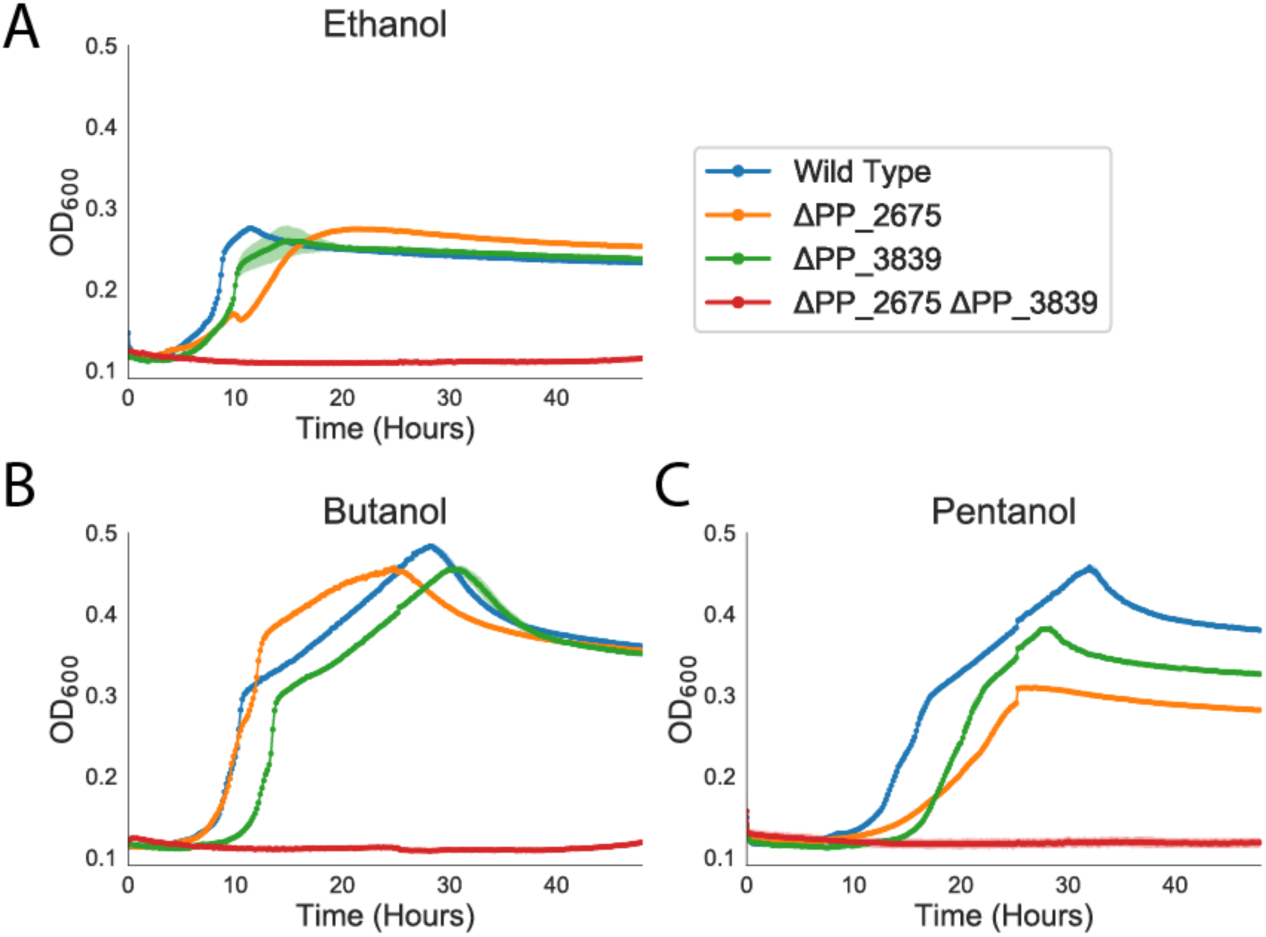
Validation of alcohol dehydrogenases involved in short chain alcohol metabolism. Growth curves of wild type (blue), ΔPP_2675 (orange), ΔPP_3839 (green), and ΔPP_2675ΔPP_3839 (red) strains of *P. putida* KT2440 on 10 mM ethanol (A), 10 mM n-butanol (B), and 10 mM n-pentanol (C). Shaded area represents 95% confidence intervals (cI), n=3.

It is ambiguous from our data and from previous work which enzymes are oxidizing the aldehyde to the corresponding carboxylic acid. As mentioned previously, both PQQ-ADHs have been biochemically shown to act on aldehydes and could catalyze the reaction, but the lack of a strong fitness phenotype for both ethanol and n-butanol suggest they are not the only enzymes capable of catalyzing this reaction. The genomically proximal aldehyde dehydrogenase *pedI* (PP_2680) showed minor fitness defects when grown on ethanol and several other alcohols (**Figure 5**, **Figure 6A**), but showed no fitness defects when libraries were grown on butanol or pentanol. Another aldehyde dehydrogenase, *aldB-I* (PP_0545), showed virtually no fitness defects when grown on any of the short chain n-alcohols tested here (**Figure 6A**). The lack of any one obvious enzyme with a distinct fitness defect supports the notion that multiple enzymes are present and able to catalyze the oxidation of these aldehydes.

While the metabolism of alcohols and their corresponding fatty acids are similar, their fitness patterns showed distinct differences. When grown on acetate, mutants in *gacS or gacA* (PP_1650 and PP_4099 - a two-component (TCS) system), *sigS* (PP_1623 - the stationary phase sigma factor sigma S), and *ptsH* (PP_0948 - a component of the sugar phosphotransferase system (PTS)) showed large and significant fitness benefits, which were not apparent when grown on ethanol (**Figure 5B**). The GacS/GacA TCS is widespread across many gram-negative bacteria, and is believed to exert transcriptional control over a wide variety of functions, sometimes in concert with a small RNA binding protein (CsrA) that exerts post-transcriptional control (56). In *Pseudomonads*, the GacA/GacS TCS has been implicated in positively controlling *sigS* expression in multiple species (57). In *P. putida* specifically, *gacS* mutations in strains engineered to produce muconic acid have resulted in higher titers (58), but disruption of the gene was also shown to completely abolish production of medium-chain length polyhydroxyalkanoates (PHAs) (59). Growth on butyrate also showed that *gacS*, *gacR*, *sigS*, and another component of the PTS (*ptsP*) had significant fitness benefits if disrupted, which was not observed when the library was grown on butanol (**Figure 5C**). Interestingly, mutants in *gacA* and *gacS* seemed to have fitness benefits when grown on either pentanol or valerate (**Figure 5D**). Further work is necessary to precisely characterize the nature of the benefits that occur when these genes are disrupted.

When grown on ethanol compared to acetate, relatively few genes not involved in the oxidation of the short chain alcohols were found to be specifically and significantly unfit; however, specific phenotypes for acetate catabolism were observed (**Figure 5B**). Mutants in PP_1635 (a two-component system response regulator), PP_1695 (variously annotated as a sodium-solute symporter, sensory box histidine kinase, or response regulator), and *tal* (PP_2168 - transaldolase) all showed fitness defects on acetate that were not observed when libraries were grown on ethanol. The high cofitness between PP_1635 and PP_1695 observed across all publicly available fitness data (*r* = 0.88) and share homology to *crbSR* systems of other bacteria where it is known to regulate acetyl-coA synthetase (60).

Much like ethanol and acetate, there were relatively few genes that showed specific fitness defects when grown on butanol that were not also observed in butyrate. However, the genes *glgB* (PP_4058 - a 1,4-alpha-glucan branching enzyme), and the co-transcribed PP_2354 and PP_2356 (annotated as a histidine kinase/response regulator (HK/RR), and histidine kinase respectively) showed specific fitness defects when grown on butyrate relative to butanol. PaperBLAST analysis of PP_2356 and PP_2354 did not reveal any publications that had explored the function of this system, and thus further work will be needed to better characterize its regulon (61). Mutants of genes encoding for three TCSs were found to be specifically unfit when grown on pentanol when compared to valerate. PP_2683 (a two component HK/RR), and *pedR1* (PP_2665 - RR) were both specifically unfit and, as previously described, are involved in the regulation of the *ped* cluster (**Figure 5D**). The gene *cbrB* (PP_4696 - sig54-dependent RR) also showed pentanol-specific defects, and is known to regulate central carbon metabolism and amino acid uptake in the *Pseudomonads* (62, 63).

### Short chain diol catabolism

Another group of industrially relevant alcohols with potential for biotechnological production are short chain diols. These compounds have broad utility ranging from plasticizers to food additives (64). As shown in **Figure 5**, most of the tested short chain diols result in significant fitness defects in *pedR1*, indicating that some of the genes involved in these metabolisms are in the PedR1 regulon. However, only 1,5-pentanediol had a strong fitness defect in *pedF*, indicating that multiple dehydrogenases may act on the shorter chain diols. Additionally, both 1,2-propanediol and 1,3-butanediol cause slight defects in mutants of the aldehyde dehydrogenase PP_0545. Although there is some ambiguity as to which enzymes initially oxidize the diols to their corresponding acids, the remaining steps in 1,2-propanediol, 1,3-butanediol, and 1,5-pentanediol catabolism are much more straightforward.

Oxidation of 1,2-propanediol yields lactate, and mutants in the L-lactate permease PP_4735 (*lldP*) have a fitness of −4.3 when grown on 1,2-propanediol. Furthermore, under this condition, mutants of the L- and D-lactate dehydrogenases PP_4736 (*lldD*) and PP_4737 (*lldE*) have fitness defects of −5.0 and −1.5, respectively. Since we provided a rac-1,2-propanediol as a substrate, this likely explains the fitness defects observed in both dehydrogenases (65, 66). Given these results, it appears that 1,2-propanediol is assimilated into central metabolism via oxidation to pyruvate (**Figure S6**).

When grown on 1,3-butanediol, two oxidations of 1,3-butanediol result in 3-hydroxybutyrate, and we observe fitness defects of −2.5 in the D-3-hydroxybutyrate dehydrogenase PP_3073 and −1.8 in the neighboring sigma-54 dependent regulator PP_3075 (67). Dehydrogenation of 3-hydroxybutyrate results in acetoacetate, and we see a fitness defect of −2.9 and −3.0 for the subunits of the predicted 3-oxoacyl-CoA transferase PP_3122-3 (*atoAB*). This enzyme likely transfers a CoA from either succinyl-CoA or acetyl-CoA in order to generate acetoacetyl-CoA. Regarding transport, mutants in the D-beta-hydroxybutyrate permease PP_3074, located in the same operon as the 3-hydroxybutyrate dehydrogenase, have a fitness defect of −0.9, while mutants in the RarD permease PP_3776 have a fitness of −1.2.

Following oxidation by the aforementioned PQQ-dependent dehydrogenases and aldehyde dehydrogenases in the periplasm, an oxidized intermediate is likely transported into the cell for the next steps in the catabolism. This is supported by the observation that mutants of the predicted dicarboxylate MFS transporter PP_1400 and its two-component regulator PP_1401-2 have strong fitness defects on both alpha-ketoglutarate and 1,5-pentanediol. Furthermore, there is a −4.7 fitness defect in mutants of the L-2-hydroxyglutarate oxidase PP_2910, which catalyzes the second step in the glutarate hydroxylation pathway of glutarate catabolism. The glutarate hydroxylase PP_2909, which catalyzes the first step of this pathway, has a much slighter negative fitness of −0.6. This is expected, because glutarate can also be catabolized through a glutaryl-CoA dehydrogenation pathway, so mutants in PP_2909 can simply divert flux through the other catabolic route (12). Mutants in PP_2910 are unable to oxidize L-2-hydroxyglutarate to alpha-ketoglutarate, and likely accumulate L-2-hydroxyglutarate as a dead-end metabolite.

1,4-butanediol catabolism has been previously studied; based on the results of expression data and adaptive laboratory evolution, Li et al. proposed three potential catabolic pathways for 1,4-butanediol, including a beta-oxidation pathway (**Figure 8**) (30). Their evolved strains had mutations in the LysR activator PP_2046 that resulted in overexpression of the beta-oxidation operon PP_2047-51 (30). Interestingly, we found that when grown on 1,4 butanediol, transposon mutants of the acyl-CoA dehydrogenase PP_2048 had significant fitness benefits and no CoA-ligase mutants showed significant fitness defects. However, a fitness defect of −1.0 in PP_0356 (malate synthase) mutants suggests that there may be flux through the beta-oxidation pathway to glycolic acid and acetyl-CoA. A possible explanation for the positive fitness of PP_2048 mutants is that the beta-oxidation pathway is suboptimal in the wild type, and it may be beneficial to divert flux through the other pathway(s). This same reasoning could also explain the absence of CoA-ligases with fitness defects; however, this also could be due to the presence of multiple CoA-ligases capable of catalyzing that step. Mutants of the 3-hydroxyacyl-CoA dehydrogenase PP_2047, a *fadB* homolog which likely catalyzes the hydration and dehydrogenation steps to produce 3-oxo-4-hydroxybutyryl-CoA, had a strong fitness defect. When PP_2047 is non-functional, 4-hydroxycrotonyl-CoA likely accumulates as a deadend metabolite resulting in decreased fitness. Li et al. also showed that deletion mutants of PP_2046 are unable to grow on 1,4-butanediol (30). Our data suggests that this is because PP_2049 appears to be the main alcohol dehydrogenase acting on either 1,4-butanediol or 4-hydroxybutyrate, and is in the operon under the control of PP_2046. Although our fitness data suggests that both the oxidation to succinate and beta-oxidation pathways occur, further work is necessary to determine if the pathway to succinyl-CoA is involved in the catabolism.

**Figure 8:**
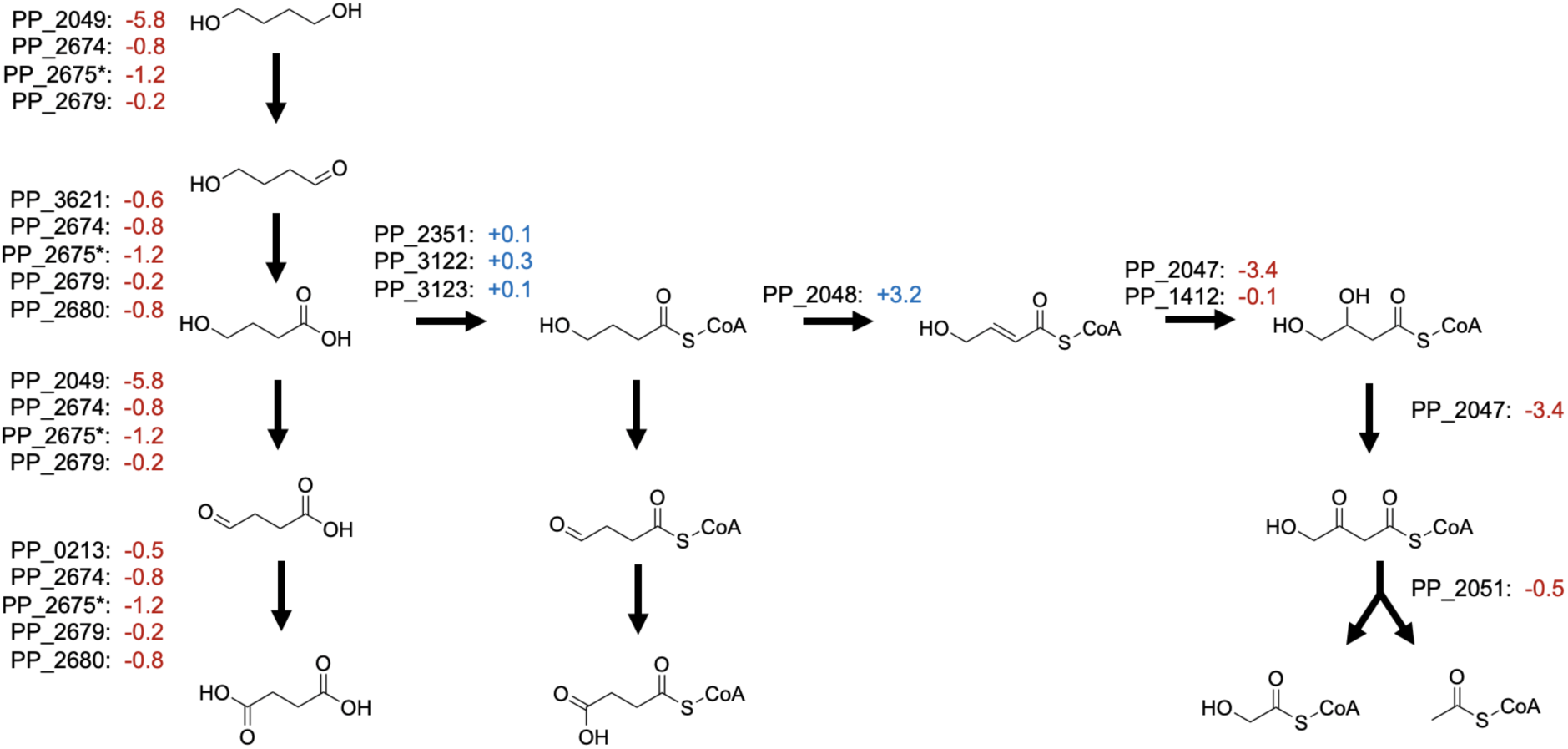
Putative routes of 1,4-butanediol catabolism in *P. putida*. Putative genes involved in catabolism of 1,4-butanediol in *P. putida*. Average fitness scores for two biological reps are shown next to each gene. The three CoA-ligases shown were proposed by Li et al.; there were no CoA-ligases that showed significant fitness defects on 1,4-butanediol. *PP_2675 (PedF) is involved in the regeneration of the PQQ cofactor predicted to be necessary for these oxidation reactions of PP_2764 (PedE) and PP_2769 (PedH).

### Branched chain alcohol metabolism

Due to their superior biofuel properties, branched chain alcohols have been targets for metabolic engineering as potential alternatives to ethanol and butanol (68). Our fitness data suggest that *pedE* and/or *pedH* oxidize 2-methyl-1-butanol to 2-methylbutyrate, which then undergoes one round of beta-oxidation to produce acetyl-CoA and propionyl-CoA (**Figure S7**). Most of the genes involved in 2-methylbutyrate beta-oxidation are located in the operon PP_2213-PP_2217. With mutants having a fitness defect of −3.2, PP_2213 appears to be the main acyl-CoA ligase acting on 2-methylbutyrate. Mutants in two predicted acyl-CoA dehydrogenases, PP_2216 and PP_0358, show fitness defects of −1.1 and −2.6, respectively. The enoyl-CoA hydratase PP_2217 has a fitness defect of −5.7 and the 3-hydroxyacyl-CoA dehydrogenase PP_2214 has a fitness defect of −5.6. Finally, the acetyl-CoA acetyltransferase appears to be PP_2215, with mutants having a fitness defect of −4.8. We also observed fitness defects of −1.8 and −1.6 in mutants of the ABC transporters, PP_5538 and PP_2667, respectively. Since 2-methylbutyrate is a known intermediate in the catabolism of isoleucine, we found that the genetic data presented here closely mirror the previous biochemical characterization of this system (69, 70).

*P. putida* can readily grow on isopentanol and isoprenol but not prenol (**Figure 9A**). All three of these alcohols have been produced in high titers in *Escherichia coli* and other bacteria because of their potential to be suitable replacements for gasoline (71, 72). RB-TnSeq data for isopentanol and isoprenol showed severe fitness defects in genes of the leucine catabolic pathway (**Figure 10**). This is unsurprising, as isopentanol can be generated from the leucine biosynthetic pathway (73). Deletion of the PP_4064-PP_4067 operon, which contains the genes that code for the conversion of isovaleryl-CoA to 3-hydroxy-3-methylglutaryl-CoA, abolished growth on both isopentanol and isoprenol (**Figure S8**). Deletion of PP_3122 (acetoacetyl CoA-transferase subunit A) also abolished growth on isopentanol, and greatly reduced growth on isoprenol (**Figure S8**). Taken together, these results validate that both of these alcohols are degraded via the leucine catabolic pathway. Transposon insertion mutants in *pedF* showed strong fitness defects on both isopentanol and isoprenol, suggesting that *pedH* (PP_2679) and *pedE* catalyze (PP_2674) the oxidation of the alcohols. Deletion mutants in *pedH* showed only a minor delay in growth compared to wild-type when grown on either isopentanol or isoprenol, while mutants in *pedE* showed a more substantial growth defect on both alcohols (**Figure 9A**). Deletion of *pedF* (PP_2675) prevented growth on both isopentanol and nearly abolished growth on isoprenol when provided as a sole carbon source in minimal media (**Figure 9A**). When wild-type *P. putida* was grown in minimal media with 10 mM glucose and 4 mM of either isopentanol, prenol, or isoprenol, each alcohol was shown to be readily degraded with concurrent observation of increasing levels of the resultant acid (**Figure 9B**). Though *P. putida* was unable to utilize prenol as a sole carbon source, it was still able to readily oxidize prenol to 3-methyl-2-butenoic acid, suggesting there is no CoA-ligase present in the cell able to activate this substrate and channel it towards leucine catabolism (**Figure 10**). When wild-type *P. putida* was grown in LB medium supplemented with 4 mM of each alcohol individually, all alcohols were completely degraded by 24 hours post-inoculation (**Figure 9C**). In *pedF* deletion mutants grown under the same conditions, the rate at which the alcohols were degraded was significantly slowed; however after 48 hours ∼50% of isopentanol, ∼75% of isoprenol, and 100% of prenol were degraded (**Figure 9C**). Uninoculated controls showed that no alcohol was lost at greater than 5% on account of evaporation (data not shown). Future efforts to produce any of these alcohols in *P. putida* will be heavily impacted by this degradation, and greater effort will need to be made to identify other enzymes involved in the oxidation of these alcohols or other metabolic pathways that consume them.

**Figure 9:**
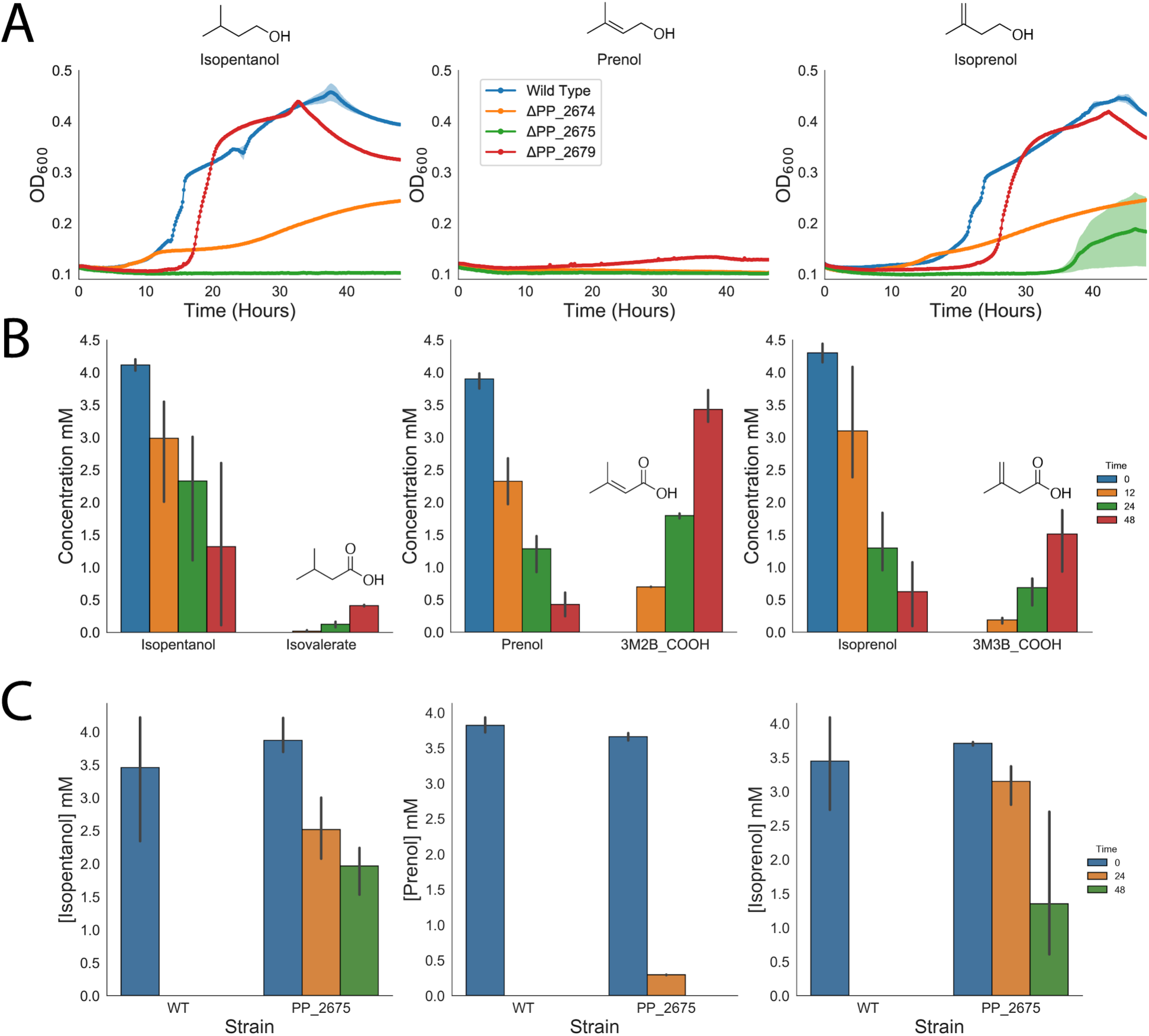
Isopentanol, Prenol, and Isoprenol consumption by *P. putida*. A) Growth curves of wild type (blue), and ΔPP_2674 (orange), ΔPP_2675 (green), and ΔPP_2679 (red) strains of *P. putida* on isopentanol (left), prenol (middle), and isoprenol (right). Structure of alcohols are shown above graphs. Shaded area represents 95% confidence intervals (cI), n=3. B) Concentrations of alcohols consumed and their corresponding carboxylic acids produced over time by wild type. Left panel shows isopentanol and isovalerate, middle panel shows prenol and 3-methyl-2-butenoic acid, and the right panel shows isoprenol and 3-methyl-3-butenoic acid. Structures of corresponding carboxylic acids derived from alcohol are shown in graphs. Error bars represent 95% cI, n=3. C) Consumption of isopentanol (left), prenol (middle), and isoprenol (right) by wild type and ΔPP_2675 *P. putida* over time. Error bars represent 95% cI, n=3.

**Figure 10:**
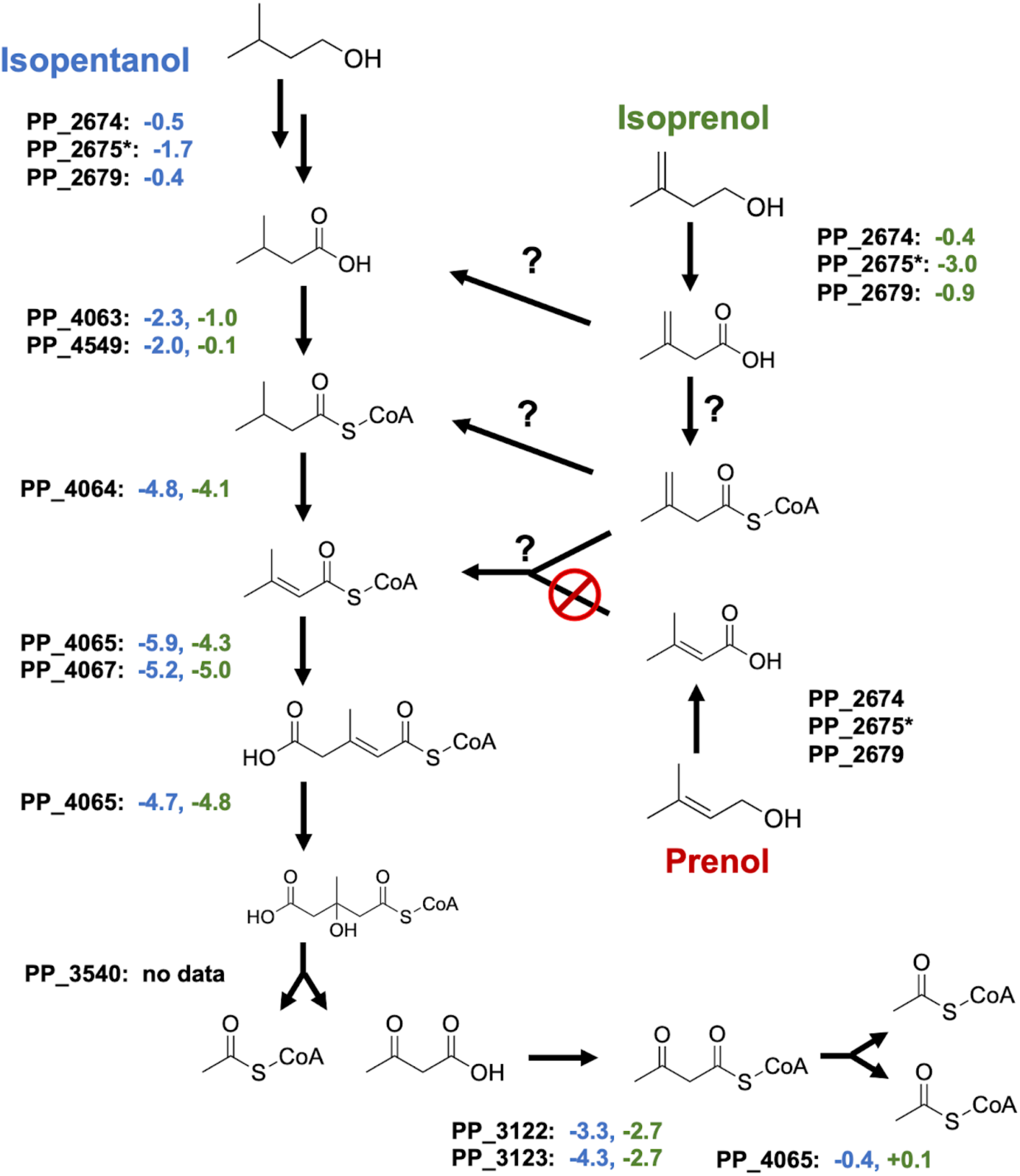
Putative routes of isopentanol and isoprenol catabolism in *P. putida*. Diagram shows the proposed pathways for the catabolism of isopentanol and isoprenol. Average fitness scores of two biological replicates for individual genes can be found next to each gene. Fitness values for isopentanol are shown in blue, while fitness values for isoprenol and shown in green. Potential reactions that would bring isoprenol into leucine catabolism are marked with a question mark.

**Figure 11:**
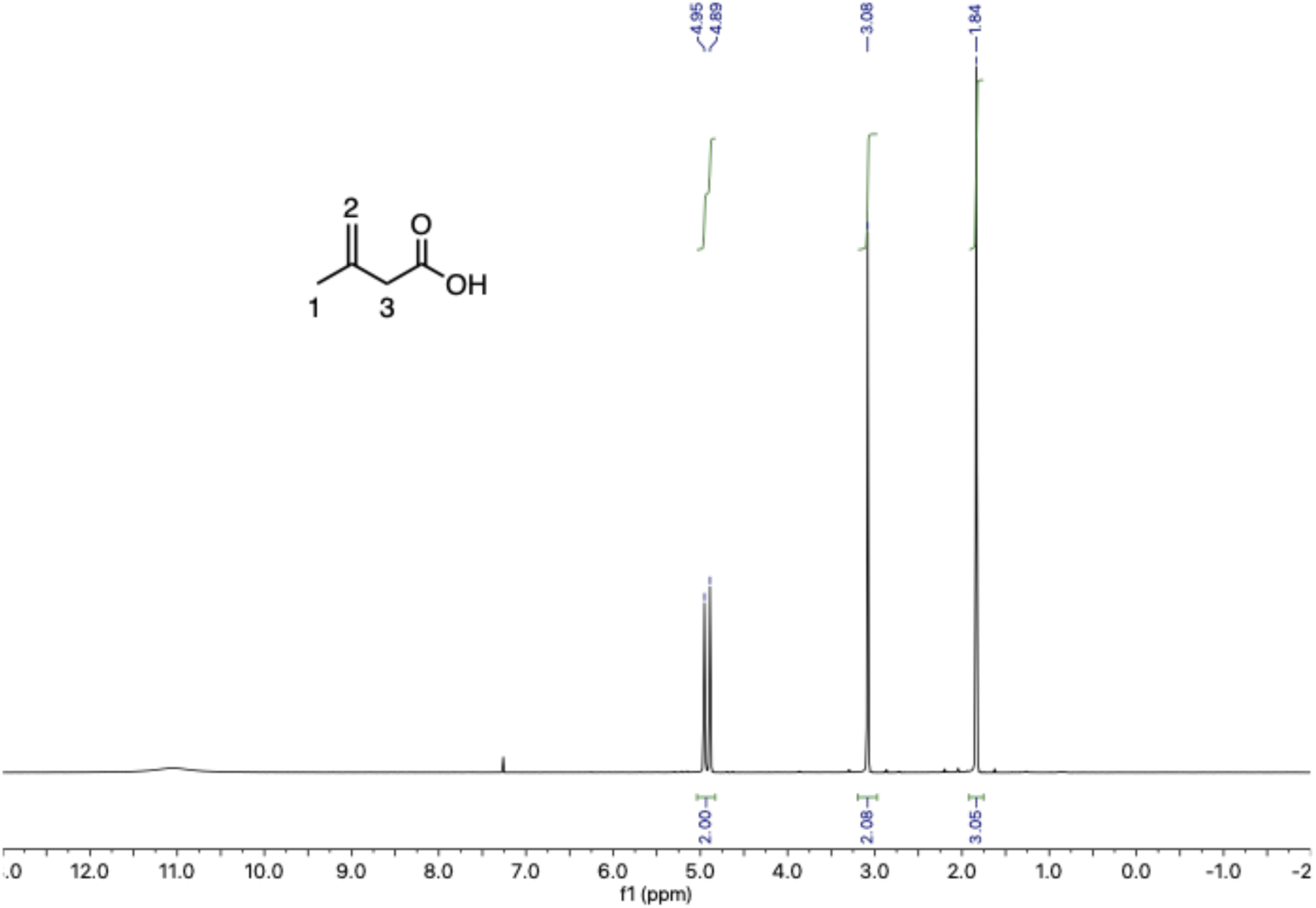
NMR validation of 3-methyl-3-butenoic acid.

One mystery that remains is how isoprenol enters into leucine catabolism. GC-MS analysis confirmed oxidation of the alcohol to 3-methyl-3-butenoic acid, but it is unclear what the next step entails. Fitness data suggests that either PP_4063 or PP_4549 may attach the CoA to isovalerate, but neither of these genes have strong phenotypes when mutant libraries are grown on isoprenol (**Figure 10**). That PP_4064 (isovaleryl-CoA dehydrogenase) shows strong negative fitness values when libraries are grown on isoprenol implies that its degradation goes through an isovaleryl-CoA intermediate, however this fitness defect may be the result of polar effects that disrupt the downstream steps (**Figure 10**). One possibility is that 3-methyl-3-butenoic acid is reduced to isovalerate in the cell; however, this seems unlikely since no isovalerate was observed via GC-MS when *P. putida* was fed isoprenol and glucose. Two other possible routes could result from the activation of 3-methyl-3-butenoic acid by an undetermined CoA-ligase. If this CoA-ligase exists, it is interesting that it would have activity on 3-methyl-3-butenoic acid but not 3-methyl-2-butenoic acid, which accumules when *P. putida* is grown in the presence of prenol. Once formed, the 3-methyl-3-butenyl-CoA could be directed into leucine catabolism via either an isomerization to 3-methylcrotonyl-CoA or a reduction to isovaleryl-CoA. Future work that leverages metabolomics to identify compounds that accumulate in leucine catabolic mutants may reveal the missing steps and help narrow the search for their enzymes.

### Future Directions

The large set of global fitness data generated in this study provide an extensive and global overview on the putative pathways of alcohol and fatty acid degradation in *P. putida*. Overall, our fitness data agree with previously published biochemical data that explored enzymes in both fatty acid and alcohol metabolism. However, there are still many questions that our data leave unanswered. Further investigation will be required to untangle and elucidate which specific enzymes are biologically relevant in the beta-oxidation of short chain fatty acids. It is likely that biochemical characterization of individual enzymes will be required to determine which of the *fad* homologs catalyze these reactions. Another intriguing question is the function of PP_0765 and PP_0766. Biochemical interrogation and mutational analysis of the DUF1302 and DUF1329 family proteins are needed to determine whether these proteins indeed function as an esterase or, as previously predicted, play some other role in outer membrane biogenesis (41). Additional work is also warranted to ascertain which of the proposed 1,4-butanediol catabolic routes the wild-type organism actually uses and determine whether the beta-oxidation pathway is indeed less preferable than the pathway to succinate.

To our knowledge, our finding that *P. putida* can consume both isopentanol and isoprenol are the first observations of this metabolism. If metabolic engineers wish to produce these chemicals in *P. putida*, these pathways will need to be removed. Critically, researchers will need to identify other enzymes that result in the oxidation of these alcohols or other routes of degradation within *P. putida*. How *P. putida* is able to utilize isoprenol, but not prenol, as a sole carbon source is metabolically intriguing. One of our proposed pathways of isoprenol catabolism requires the existence of a CoA-ligase that shows surprising specificity towards 3-methyl-3-butenoic acid with little to no activity on 3-methyl-2-butenoic acid. More work should be done to leverage other omics-levels techniques to try to identify this hypothetical enzyme and biochemically verify its activity. Finally, this data set as a whole will likely strengthen the assumptions made by genome-scale metabolic models. Previous models of *P. putida* metabolism have incorporated RB-TnSeq data to improve their predictions (17). This work nearly doubles the number of available RB-TnSeq datasets in *P. putida* that are publicly available and will likely contribute greatly to further model refinement. Ultimately, large strides in our understanding of *P. putida* metabolism leveraging functional genomic approaches will provide the foundation for improved metabolic engineering efforts in the future.

## Methods

### Media, chemicals, and culture conditions

General *E. coli* cultures were grown in lysogeny broth (LB) Miller medium (BD Biosciences, USA) at 37 °C while *P. putida* was grown at 30 °C. When indicated, *P. putida* and *E. coli* were grown on modified MOPS minimal medium, which is comprised of 32.5 µM CaCl_2_, 0.29 mM K_2_SO_4_, 1.32 mM K_2_HPO_4_, 8 µM FeCl_2_, 40 mM MOPS, 4 mM tricine, 0.01 mM FeSO_4_, 9.52 mM NH_4_Cl, 0.52 mM MgCl_2_, 50 mM NaCl, 0.03 µM (NH_4_)_6_Mo_7_O_24_, 4 µM H_3_BO_3_, 0.3 µM CoCl_2_, 0.1 µM CuSO_4_, 0.8 µM MnCl_2_, and 0.1 µM ZnSO_4_ (74). Cultures were supplemented with kanamycin (50 mg/L, Sigma Aldrich, USA), gentamicin (30 mg/L, Fisher Scientific, USA), or carbenicillin (100mg/L, Sigma Aldrich, USA), when indicated. All other compounds were purchased through Sigma Aldrich (Sigma Aldrich, USA). 3-methyl-3-butenoic acid was not available commercially and required synthesis which is described below.

### Strains and plasmids

All bacterial strains and plasmids used in this work are listed in Table 1. All strains and plasmids created in this work are available through the public instance of the JBEI registry. (public-registry.jbei.org/folders/456). All plasmids were designed using Device Editor and Vector Editor software, while all primers used for the construction of plasmids were designed using j5 software (75–77). Plasmids were assembled via Gibson Assembly using standard protocols (78), or Golden Gate Assembly using standard protocols (79). Plasmids were routinely isolated using the Qiaprep Spin Miniprep kit (Qiagen, USA), and all primers were purchased from Integrated DNA Technologies (IDT, Coralville, IA). Construction of *P. putida* deletion mutants was performed as described previously (18).

**Table 1:** Strains and plasmids used in this study.

| Strain | Description | Reference |
| --- | --- | --- |
| *E. coli* XL1 Blue |  | Agilent |
| *P. putida* KT2440 | Wild-Type | ATCC 47054 |
| *P. putida* ΔPP_2674 | Strain with complete internal in-frame deletion of PP_2674 | This study |
| *P. putida* ΔPP_2675 | Strain with complete internal in-frame deletion of PP_2675 | This study |
| *P. putida* ΔPP_2679 | Strain with complete internal in-frame deletion of PP_2679 | This study |
| *P. putida* ΔPP_3839 | Strain with complete internal in-frame deletion of PP_3839 | This study |
| *P. putida* ΔPP_2675 ΔPP_3839 | A double knockout of PP_2675 and PP_3839 | This study |
| *P. putida* ΔPP_4064-PP_4067 | Strain with complete internal in-frame deletion of the PP_4064-4067 operon | This study |
| *P. putida* ΔPP_3122 | Strain with complete internal in-frame deletion of PP_3122 | This study |
| **Plasmids** |  |  |
| pMQ30 | Suicide vector for allelic replace Gm<sup>r</sup>, SacB | (80) |
| pMQ30 ΔPP_2674 | pMQ30 derivative harboring 1kb flanking regions of PP_2674 | This study |
| pMQ30 ΔPP_2675 | pMQ30 derivative harboring 1kb flanking regions of PP_2675 | This study |
| pMQ30 ΔPP_2679 | pMQ30 derivative harboring 1kb flanking regions of PP_2679 | This study |
| pMQ30 ΔPP_3839 | pMQ30 derivative harboring 1kb flanking regions of PP_3839 | This study |
| pMQ30 ΔPP_4064-PP_4067 | pMQ30 derivative harboring 1kb flanking regions of PP_4064 and PP_4067 | This study |
| pMQ30 ΔPP_3122 | pMQ30 derivative harboring 1kb flanking regions of PP_3122 | This study |

### Plate-based growth assays

Growth studies of bacterial strains were conducted using microplate reader kinetic assays as described previously (81). Overnight cultures were inoculated into 10 mL of LB medium from single colonies, and grown at 30 °C. These cultures were then washed twice with MOPS minimal media without any added carbon and diluted 1:100 into 500 μL of MOPS medium with 10 mM of a carbon source in 48-well plates (Falcon, 353072). Plates were sealed with a gas-permeable microplate adhesive film (VWR, USA), and then optical density and fluorescence were monitored for 48 hours in an Biotek Synergy 4 plate reader (BioTek, USA) at 30 °C with fast continuous shaking. Optical density was measured at 600 nm.

### RB-TnSeq

RB-TnSeq experiments utilized *P. putida* library JBEI-1 which has been described previously with slight modification(18). Libraries of JBEI-1 were thawed on ice, diluted into 25 mL of LB medium with kanamycin and then grown to an OD_600_ of 0.5 at 30 °C at which point three 1-mL aliquots were removed, pelleted, and stored at −80 °C. Libraries were then washed once in MOPS minimal medium with no carbon source, and then diluted 1:50 in MOPS minimal medium with 10 mM of each carbon source tested. Cells were grown in 10 mL of medium in test tubes at 30 °C shaking at 200 rpm. One 500-μL aliquot was pelleted, and stored at −80 °C until BarSeq analysis, which was performed as previously described (19, 40). The fitness of a strain is defined here as the normalized log_2_ ratio of barcode reads in the experimental sample to barcode reads in the time zero sample. The fitness of a gene is defined here as the weighted average of the strain fitness for insertions in the central 10% to 90% of the gene. The gene fitness values are normalized such that the typical gene has a fitness of zero. The primary statistic *t* value represents the form of fitness divided by the estimated variance across different mutants of the same gene. Statistic *t* values of >|4| were considered significant. A more detailed explanation of calculating fitness scores can be found in Wetmore et al. (40). All experiments described here passed testing using the quality metrics described previously unless noted otherwise (40). All experiments were conducted in biological duplicate, and all fitness data are publically available at http://fit.genomics.lbl.gov.

### GC-MS and GC-FID Analysis of Branched Alcohol Consumption

To examine the oxidation of isopentanol, prenol, and isoprenol to their corresponding acids 10mL of MOPS minimal medium supplemented with 10 mM glucose and 4mM of one of each alcohol added were inoculated with a 1:100 dilution of overnight *P. putida* culture and incubated at 30 °C with 200 rpm shaking. At 0, 12, 24, and 48-hours post-inoculation 200 μL of media were sampled and stored at −80 °C. Alcohols and fatty acids were extracted by acidifying media with 10 μL of 10N HCl, followed by addition of an 200 μL of ethyl-acetate. To detect alcohols and their corresponding carboxylic acids via GC-MS an Agilent 6890 system equipped with a DB-5ms column (30-m×0.25 mm×0.25 µm) and an Agilent 5973 MS were used. Helium (constant flow 1 mL/min) was used as the carrier gas. The temperature of the injector was 250 °C and the following temperature program was applied: 40 °C for 2 min, increase of 10 °C/min to 100 °C then increase of 35 °C/min to 300 °C, temperature was then held at 300 °C for 1 min. Authentic standards were used to quantify analytes. Determination of isopentanol, prenol, and isoprenol consumption was conducted in 10mL LB medium with 4mM of either alcohol added. Cultures were inoculated with a 1:100 dilution of overnight *P. putida* culture and incubated at 30 °C with 200 rpm shaking. At 0, 24, and 48 hours post-inoculation 200 μL of media were sampled and stored at −80 °C. The remaining concentration of each alcohol was determined by GC-FID as previously described (82).

### Synthesis of 3-Methyl-3-Butenoic Acid

To a 25-mL round bottom flask was added chromium(VI) oxide (0.69 g, 6.9 mmol) and distilled water (1 mL). The reaction mixture was then cooled to 0 °C before concentrated sulfuric acid (0.6 mL, 10.5 mmol) was added dropwise, thus forming Jones reagent. The solution of Jones reagent was then diluted to a total volume of 5 mL with distilled water. To a stirred solution of 3-methyl-3-buten-1-ol (0.59 g, 6.9 mmol) in acetone (7 ml) was added dropwise the Jones reagent at 0 °C. After being stirred for 8 h at room temperature, the mixture was quenched with ethanol. The mixture was then diluted with water, and acetone was evaporated *in vacuo*. The residue was extracted with DCM, and organic layers were combined and washed three times with saturated *aq.* NaHCO_3_ solution. The aqueous phase was acidified with a 2 M *aq.* HCl solution to pH 2-3, which was then extracted again with DCM. The extract was successively washed with water and brine, dried over MgSO_4_, and concentrated *in vacuo*. The residue was distilled (90 °C, 100 mTorr) to yield 3-methyl-3-butenoic acid as a clear oil. ^1^H NMR (300 MHz, Chloroform-*d*) δ 4.92 (d, *J* = 19.1 Hz, 2H), 3.08 (s, 2H), 1.84 (s, 3H) (**Figure 10**).

### Bioinformatic Analyses

PaperBLAST was routinely used to search for literature on proteins of interest and related homologs (61). All statistical analyses were carried out using either the Python Scipy or Numpy libraries (83, 84). For the phylogenetic reconstructions, the best amino acid substitution model was selected using ModelFinder as implemented on IQ-tree (85) phylogenetic trees were constructed using IQ-tree, nodes were supported with 10,000 bootstrap replicates. The final tree figures were edited using FigTree v1.4.3 (http://tree.bio.ed.ac.uk/software/figtree/). Orthologous syntenic regions were identified with CORASON-BGC (86) and manually colored and annotated.

## Acknowledgements

We would like to thank Morgan Price for assistance in analyzing RB-TnSeq data. The laboratory of LMB is partially funded by the Deutsche Forschungsgemeinschaft (DFG, German Research Foundation) under Germany’s Excellence Strategy within the Cluster of Excellence FSC 2186 ‘The Fuel Science Center’. This work was part of the DOE Joint BioEnergy Institute (https://www.jbei.org) supported by the U. S. Department of Energy, Office of Science, Office of Biological and Environmental Research, supported by the U.S. Department of Energy, Energy Efficiency and Renewable Energy, Bioenergy Technologies Office, through contract DE-AC02-05CH11231 between Lawrence Berkeley National Laboratory and the U.S. Department of Energy. The views and opinions of the authors expressed herein do not necessarily state or reflect those of the United States Government or any agency thereof. Neither the United States Government nor any agency thereof, nor any of their employees, makes any warranty, expressed or implied, or assumes any legal liability or responsibility for the accuracy, completeness, or usefulness of any information, apparatus, product, or process disclosed, or represents that its use would not infringe privately owned rights. The United States Government retains and the publisher, by accepting the article for publication, acknowledges that the United States Government retains a nonexclusive, paid-up, irrevocable, worldwide license to publish or reproduce the published form of this manuscript, or allow others to do so, for United States Government purposes. The Department of Energy will provide public access to these results of federally sponsored research in accordance with the DOE Public Access Plan (http://energy.gov/downloads/doe-public-access-plan).

## Contributions

Conceptualization, M.G.T., M.R.I., A.N.P.; Methodology, M.G.T., M.R.I., A.N.P., J.M.B, P.C.M., A.M.D.; Investigation, M.G.T., M.R.I., A.N.P, M.S., W.A.S., C.B.E., P.C.M., J.M.B., Y.L., R.W.H., C.A.A, R.N.K, P.L.; Writing – Original Draft, M.G.T., M.R.I., A.N.P.; Writing – Review and Editing, All authors.; Resources and supervision, L.M.B., A.M., A.M.D., P.M.S, J.D.K.

M.G.T., M.R.I., and A.N.P. contributed equally to this work. Author order was determined by the outcome of a MarioKart 64 tournament.

## Competing Interests

J.D.K. has financial interests in Amyris, Lygos, Demetrix, Napigen, Maple Bio, and Apertor Labs. C.B.E has a financial interest in Perlumi Chemicals.

## Supplemental Figures

**Figure S1:**
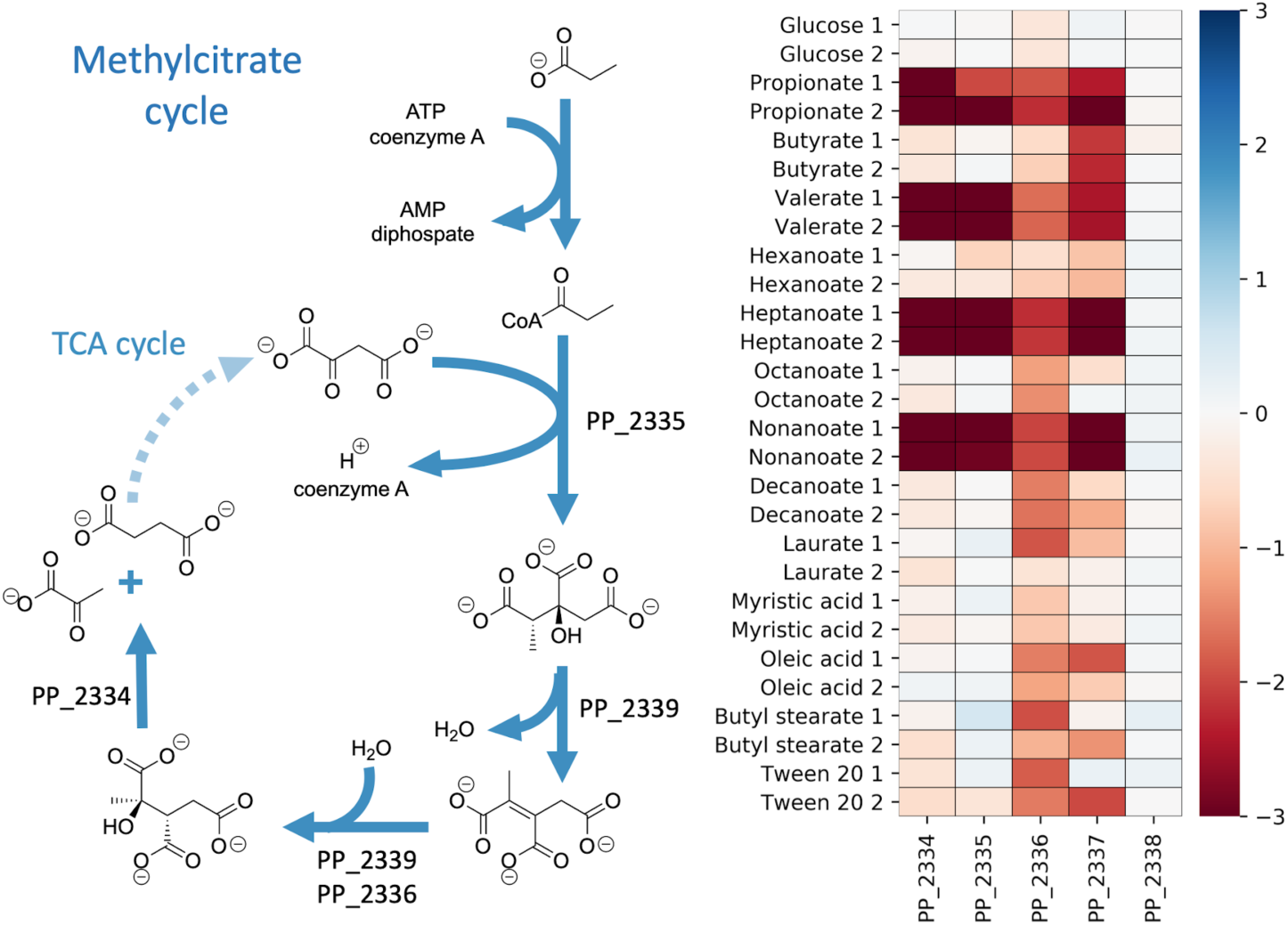
Fitness scores for genes catalyzing the methylcitrate cycle of *P. putida* KT2440. Metabolic pathway for the MCC of *P. putida* KT2440. Heatmap to the right shows fitness scores for MCC genes when grown on fatty acids or glucose. PP_2337 is predicted to function as an aconitate isomerase not shown in the depiction (left).

**Figure S2:**
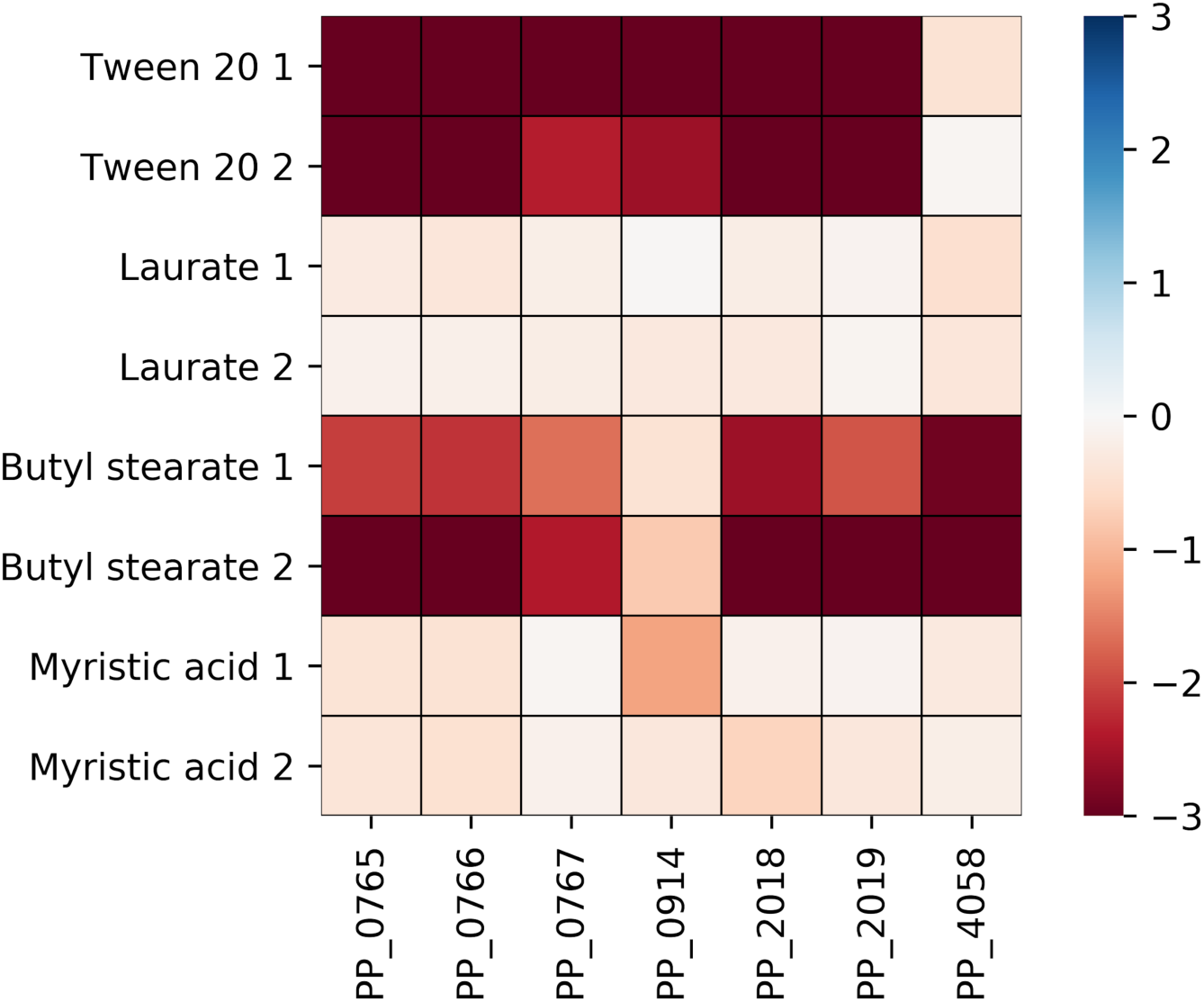
Genes in *P. putida* specifically defective for growth on long chain fatty esters compared to long chain fatty acids. Heatmap shows fitness values of genes that were found to be specifically defective on fatty esters. Conditions shown are the two fatty esters Tween 20 and butyl stearate, as well as long chain fatty acids laurate and myristate.

**Figure S3:**
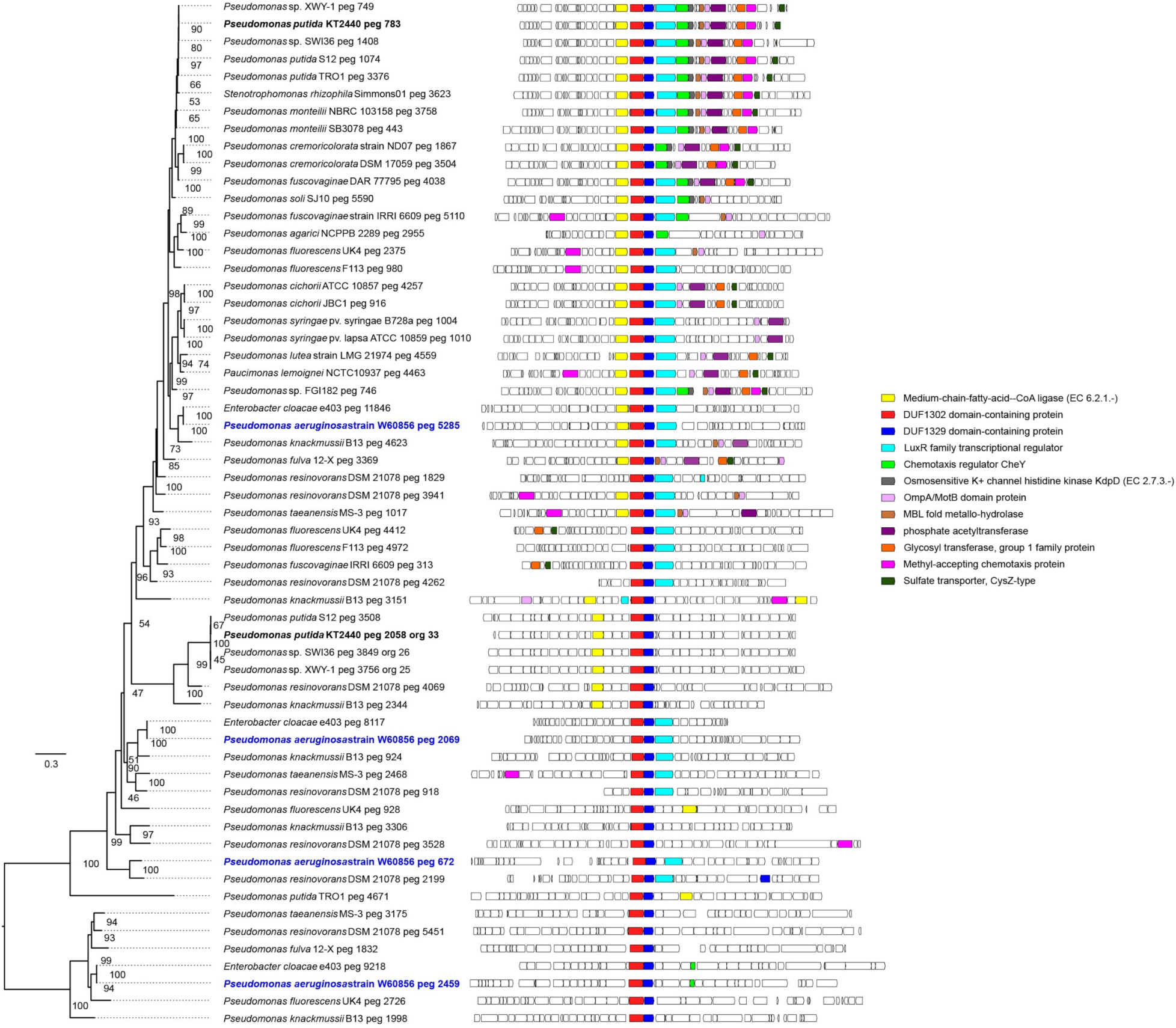
Genomic context analysis of PP_0765/PP_0766 homologs across related *Pseudomonas* species. The genomic contexts of PP_0765/PP_0766 orthologs across multiple related *Pseudomonas* species are shown. Phylogenetic tree shows relatedness of the conserved gene pair. Genes that are commonly found near PP_0765/PP_0766 are colored, medium chain fatty acid coA-ligases are shown in yellow.

**Figure S4:**
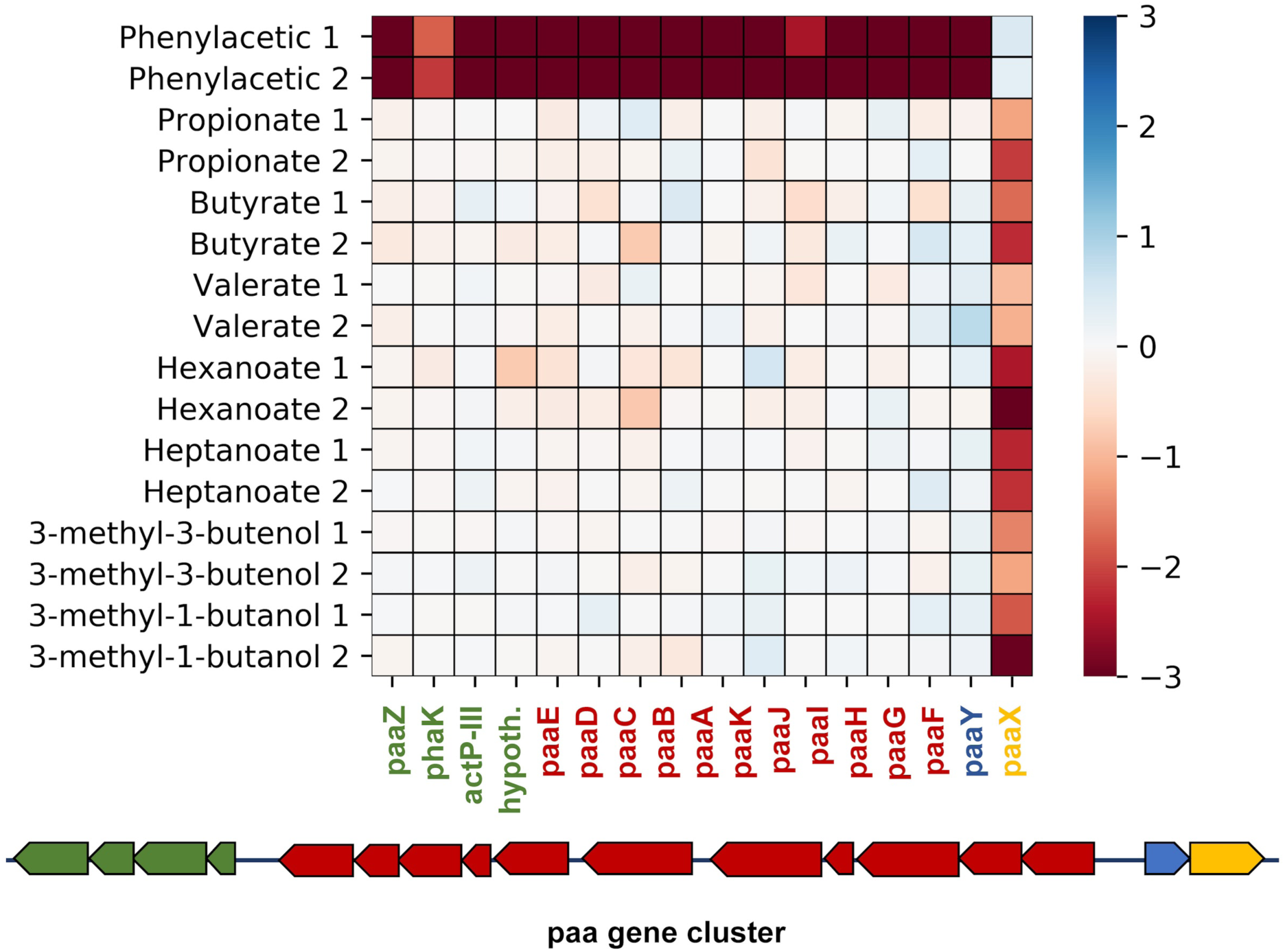
Detrimental fitness effects on short chain fatty acid catabolism by disrupting *paaX.* Heatmap shows fitness values for the *paa* gene cluster when grown on short chain fatty acids, isoprenol, isopentanol, and phenylacetic acid.

**Figure S5:**
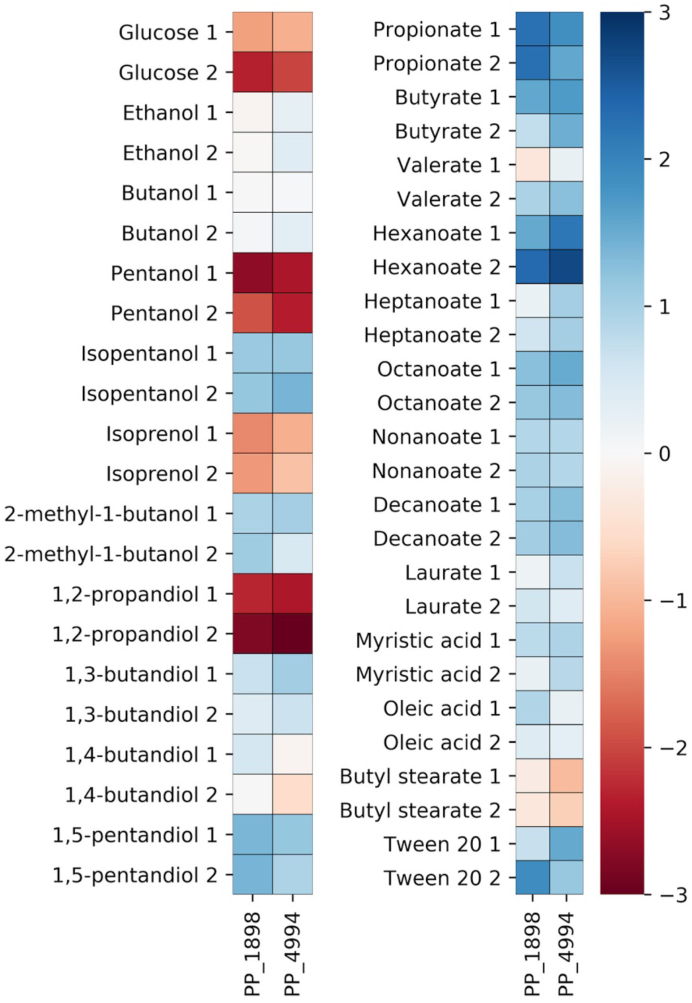
Fitness profiles of tonB siderophore transporter when grown on fatty acids or alcohols. Heatmap shows fitness scores for PP_1898, and PP_4494 on all conditions tested in this work.

**Figure S6:**
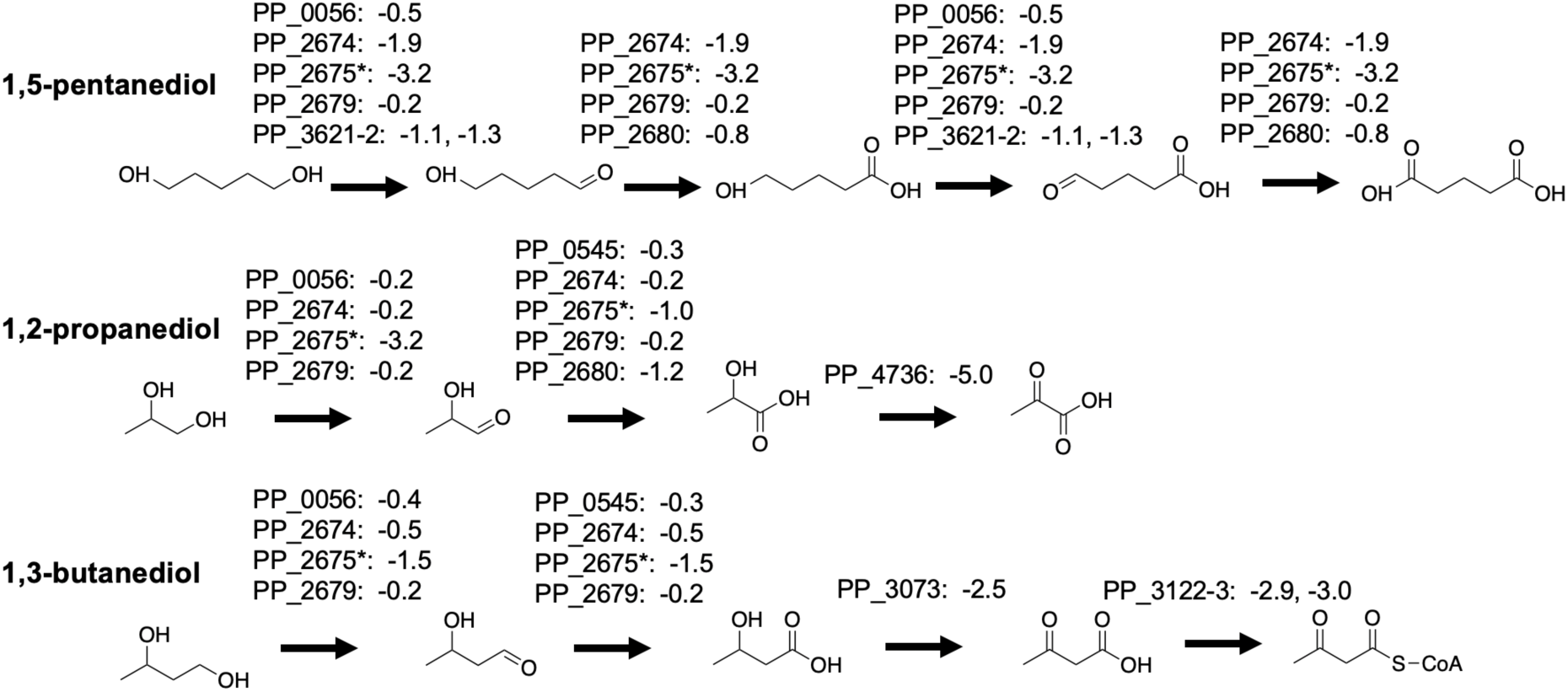
Putative catabolic pathways for 1,5-pentanediol, 1,2-propanediol, and 1,3-butanediol. Fitness scores for two biological replicates of genes proposed to code for responsible enzymes can be found next to genes.

**Figure S7:**
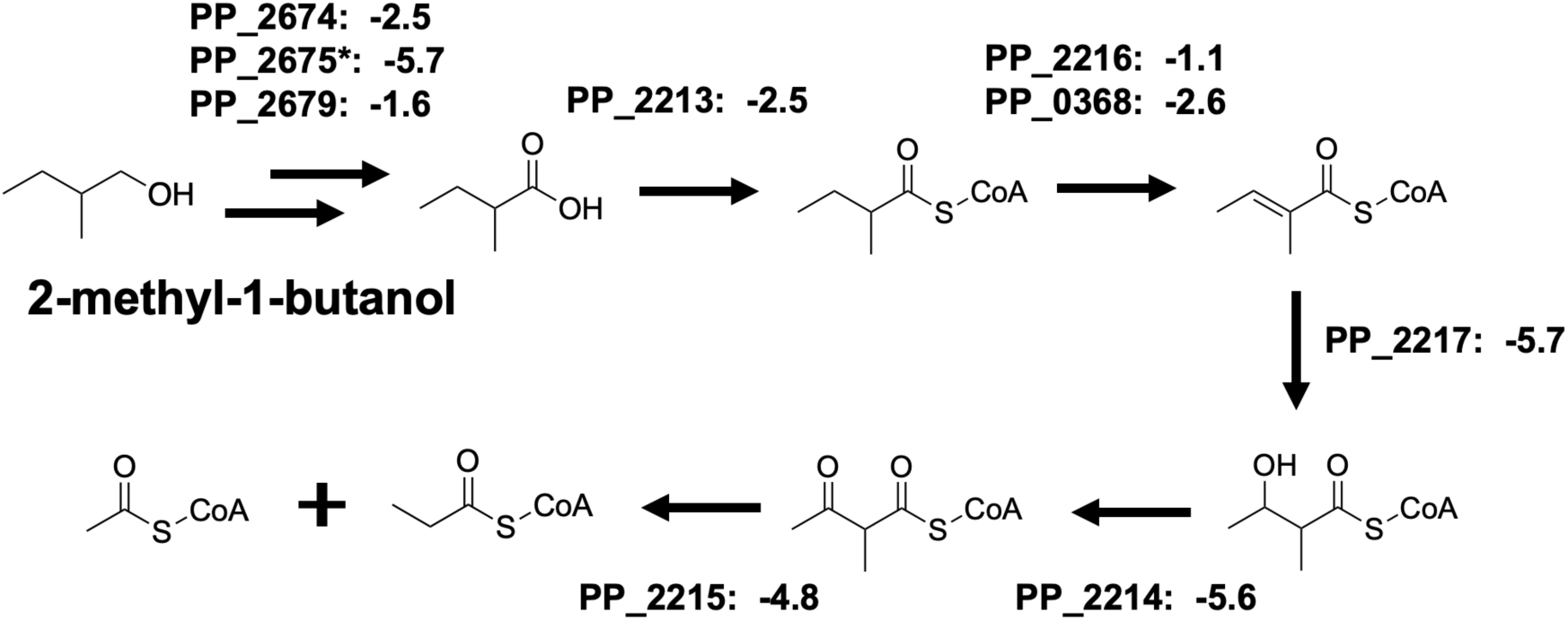
Putative catabolic pathway for 2-methyl-1-butanol. Fitness scores for two biological replicates of genes proposed to code for responsible enzymes can be found next to genes.

**Figure S8:**
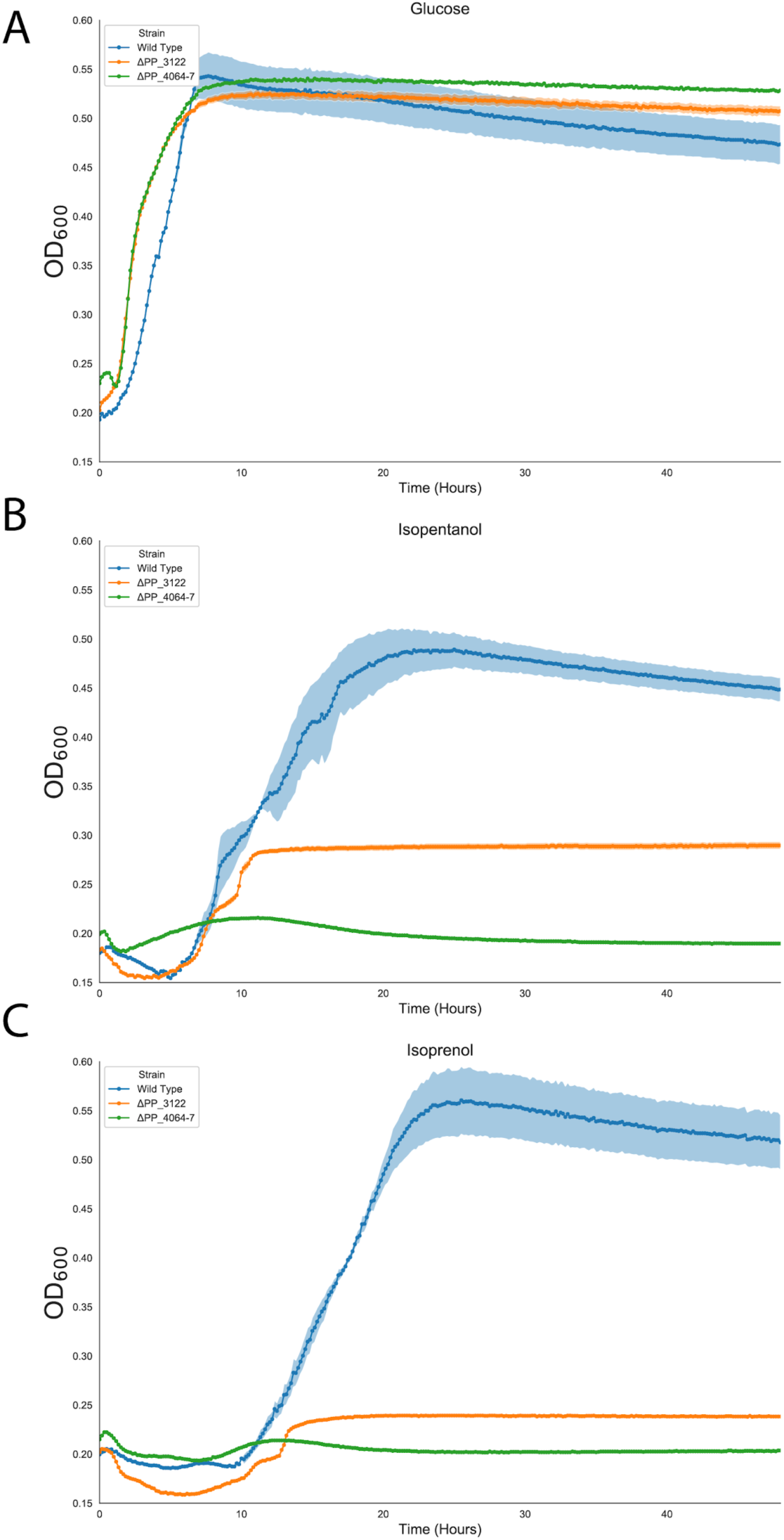
Growth of leucine catabolism deletion mutants on branched chain alcohols. Growth curves of wild-type (blue), ΔPP_3122 (orange), and ΔPP_4064-4067 (green) strains of *P. putida* on glucose (A), isopentanol (B), and isoprenol (C). Shaded area represents 95% cI, n=4.

## Bibliography

1. Park M, Chen Y, Thompson M, Benites VT, Fong B, Petzold CJ, Baidoo EEK, Gladden JM, Adams PD, Keasling JD, Simmons BA, Singer SW. 2020. Response of *Pseudomonas putida* to Complex, Aromatic-Rich Fractions from Biomass. ChemSusChem 13:1–14.

2. Belda E, van Heck RGA, José Lopez-Sanchez M, Cruveiller S, Barbe V, Fraser C, Klenk H-P, Petersen J, Morgat A, Nikel PI, Vallenet D, Rouy Z, Sekowska A, Martins Dos Santos VAP, de Lorenzo V, Danchin A, Médigue C. 2016. The revisited genome of Pseudomonas putida KT2440 enlightens its value as a robust metabolic chassis. Environ Microbiol 18:3403–3424.

3. Tiso T, Narancic T, Wei R, Pollet E, Beagan N, Schröder K, Honak A, Jiang M, Kenny ST, Wierckx N, Perrin R, Avérous L, Zimmermann W, O’Connor K, Blank LM. 2020. Bio-upcycling of polyethylene terephthalate. BioRxiv.

4. Aparicio T, de Lorenzo V, Martínez-García E. 2019. CRISPR/Cas9-enhanced ssDNA recombineering for Pseudomonas putida. Microb Biotechnol 12:1076–1089.

5. Aparicio T, de Lorenzo V, Martínez-García E. 2018. CRISPR/Cas9-Based Counterselection Boosts Recombineering Efficiency in Pseudomonas putida. Biotechnol J 13:e1700161.

6. Cook TB, Rand JM, Nurani W, Courtney DK, Liu SA, Pfleger BF. 2018. Genetic tools for reliable gene expression and recombineering in Pseudomonas putida. J Ind Microbiol Biotechnol 45:517–527.

7. Banerjee D, Eng TT, Lau AK, Wang B, Sasaki Y, Herbert RA, Chen Y, Liu Y, Prahl J-P, Singan VR, Tanjore D, Petzold CJ, Keasling JD, Mukhopadhyay A. 2020. Genome-scale metabolic rewiring to achieve predictable titers rates and yield of a non-native product at scale. BioRxiv.

8. Blank LM, Ionidis G, Ebert BE, Bühler B, Schmid A. 2008. Metabolic response of Pseudomonas putida during redox biocatalysis in the presence of a second octanol phase. FEBS J 275:5173–5190.

9. Ebert BE, Kurth F, Grund M, Blank LM, Schmid A. 2011. Response of Pseudomonas putida KT2440 to increased NADH and ATP demand. Appl Environ Microbiol 77:6597–6605.

10. Thompson MG, Valencia LE, Blake-Hedges JM, Cruz-Morales P, Velasquez AE, Pearson AN, Sermeno LN, Sharpless WA, Benites VT, Chen Y, Baidoo EEK, Petzold CJ, Deutschbauer AM, Keasling JD. 2019. Omics-driven identification and elimination of valerolactam catabolism in Pseudomonas putida KT2440 for increased product titer. Metab Eng Commun 9:e00098.

11. Incha MR, Thompson MG, Blake-Hedges JM, Liu Y, Pearson AN, Schmidt M, Gin JW, Petzold CJ, Deutschbauer AM, Keasling JD. 2020. Leveraging host metabolism for bisdemethoxycurcumin production in Pseudomonas putida. Metab Eng Commun 10:e00119.

12. Zhang M, Gao C, Guo X, Guo S, Kang Z, Xiao D, Yan J, Tao F, Zhang W, Dong W, Liu P, Yang C, Ma C, Xu P. 2018. Increased glutarate production by blocking the glutaryl-CoA dehydrogenation pathway and a catabolic pathway involving L-2-hydroxyglutarate. Nat Commun 9:2114.

13. Dong J, Chen Y, Benites VT, Baidoo EEK, Petzold CJ, Beller HR, Eudes A, Scheller HV, Adams PD, Mukhopadhyay A, Simmons BA, Singer SW. 2019. Methyl ketone production by Pseudomonas putida is enhanced by plant-derived amino acids. Biotechnol Bioeng 116:1909–1922.

14. Tiso T, Zauter R, Tulke H, Leuchtle B, Li W-J, Behrens B, Wittgens A, Rosenau F, Hayen H, Blank LM. 2017. Designer rhamnolipids by reduction of congener diversity: production and characterization. Microb Cell Fact 16:225.

15. Kohlstedt M, Starck S, Barton N, Stolzenberger J, Selzer M, Mehlmann K, Schneider R, Pleissner D, Rinkel J, Dickschat JS, Venus J, B J H van Duuren J, Wittmann C. 2018. From lignin to nylon: Cascaded chemical and biochemical conversion using metabolically engineered Pseudomonas putida. Metab Eng 47:279– 293.

16. Loeschcke A, Thies S. 2020. Engineering of natural product biosynthesis in Pseudomonas putida. Curr Opin Biotechnol 65:213–224.

17. Nogales J, Mueller J, Gudmundsson S, Canalejo FJ, Duque E, Monk J, Feist AM, Ramos JL, Niu W, Palsson BO. 2020. High-quality genome-scale metabolic modelling of Pseudomonas putida highlights its broad metabolic capabilities. Environ Microbiol 22:255–269.

18. Thompson MG, Blake-Hedges JM, Cruz-Morales P, Barajas JF, Curran SC, Eiben CB, Harris NC, Benites VT, Gin JW, Sharpless WA, Twigg FF, Skyrud W, Krishna RN, Pereira JH, Baidoo EEK, Petzold CJ, Adams PD, Arkin AP, Deutschbauer AM, Keasling JD. 2019. Massively Parallel Fitness Profiling Reveals Multiple Novel Enzymes in Pseudomonas putida Lysine Metabolism. MBio 10.

19. Rand JM, Pisithkul T, Clark RL, Thiede JM, Mehrer CR, Agnew DE, Campbell CE, Markley AL, Price MN, Ray J, Wetmore KM, Suh Y, Arkin AP, Deutschbauer AM, Amador-Noguez D, Pfleger BF. 2017. A metabolic pathway for catabolizing levulinic acid in bacteria. Nat Microbiol 2:1624–1634.

20. Price MN, Ray J, Iavarone AT, Carlson HK, Ryan EM, Malmstrom RR, Arkin AP, Deutschbauer AM. 2019. Oxidative pathways of deoxyribose and deoxyribonate catabolism. mSystems 4.

21. Zhu Z, Hu Y, Teixeira PG, Pereira R, Chen Y, Siewers V, Nielsen J. 2020. Multidimensional engineering of Saccharomyces cerevisiae for efficient synthesis of medium-chain fatty acids. Nat Catal 3:64–74.

22. Luo X, Reiter MA, d’Espaux L, Wong J, Denby CM, Lechner A, Zhang Y, Grzybowski AT, Harth S, Lin W, Lee H, Yu C, Shin J, Deng K, Benites VT, Wang G, Baidoo EEK, Chen Y, Dev I, Petzold CJ, Keasling JD. 2019. Complete biosynthesis of cannabinoids and their unnatural analogues in yeast. Nature 567:123–126.

23. Mehrer CR, Incha MR, Politz MC, Pfleger BF. 2018. Anaerobic production of medium-chain fatty alcohols via a β-reduction pathway. Metab Eng 48:63–71.

24. Karp PD, Billington R, Caspi R, Fulcher CA, Latendresse M, Kothari A, Keseler IM, Krummenacker M, Midford PE, Ong Q, Ong WK, Paley SM, Subhraveti P. 2019. The BioCyc collection of microbial genomes and metabolic pathways. Brief Bioinformatics 20:1085–1093.

25. Wehrmann M, Billard P, Martin-Meriadec A, Zegeye A, Klebensberger J. 2017. Functional Role of Lanthanides in Enzymatic Activity and Transcriptional Regulation of Pyrroloquinoline Quinone-Dependent Alcohol Dehydrogenases in Pseudomonas putida KT2440. MBio 8.

26. Wehrmann M, Toussaint M, Pfannstiel J, Billard P, Klebensberger J. 2020. The Cellular Response to Lanthanum Is Substrate Specific and Reveals a Novel Route for Glycerol Metabolism in Pseudomonas putida KT2440. MBio 11.

27. Nikel PI, de Lorenzo V. 2014. Robustness of Pseudomonas putida KT2440 as a host for ethanol biosynthesis. N Biotechnol 31:562–571.

28. Cuenca M del S, Roca A, Molina-Santiago C, Duque E, Armengaud J, Gómez-Garcia MR, Ramos JL. 2016. Understanding butanol tolerance and assimilation in Pseudomonas putida BIRD-1: an integrated omics approach. Microb Biotechnol 9:100–115.

29. Simon O, Klebensberger J, Mükschel B, Klaiber I, Graf N, Altenbuchner J, Huber A, Hauer B, Pfannstiel J. 2015. Analysis of the molecular response of Pseudomonas putida KT2440 to the next-generation biofuel n-butanol. J Proteomics 122:11–25.

30. Li W-J, Narancic T, Kenny ST, Niehoff P-J, O’Connor K, Blank LM, Wierckx N. 2020. Unraveling 1,4-Butanediol Metabolism in Pseudomonas putida KT2440. Front Microbiol 11:382.

31. Basler G, Thompson M, Tullman-Ercek D, Keasling J. 2018. A Pseudomonas putida efflux pump acts on short-chain alcohols. Biotechnol Biofuels 11:136.

32. Nikel PI, Chavarría M, Danchin A, de Lorenzo V. 2016. From dirt to industrial applications: Pseudomonas putida as a Synthetic Biology chassis for hosting harsh biochemical reactions. Curr Opin Chem Biol 34:20–29.

33. Burlage RS, Hooper SW, Sayler GS. 1989. The TOL (pWW0) catabolic plasmid. Appl Environ Microbiol 55:1323–1328.

34. Nozzi NE, Desai SH, Case AE, Atsumi S. 2014. Metabolic engineering for higher alcohol production. Metab Eng 25:174–182.

35. Yan Q, Pfleger BF. 2020. Revisiting metabolic engineering strategies for microbial synthesis of oleochemicals. Metab Eng 58:35–46.

36. Pfleger BF, Gossing M, Nielsen J. 2015. Metabolic engineering strategies for microbial synthesis of oleochemicals. Metab Eng 29:1–11.

37. Kukurugya MA, Mendonca CM, Solhtalab M, Wilkes RA, Thannhauser TW, Aristilde L. 2019. Multi-omics analysis unravels a segregated metabolic flux network that tunes co-utilization of sugar and aromatic carbons in Pseudomonas putida. J Biol Chem 294:8464–8479.

38. Nikel PI, Fuhrer T, Chavarria M, Sanchez-Pascuala A, Sauer U, de Lorenzo V. 2020. Redox stress reshapes carbon fluxes of Pseudomonas putida for cytosolic glucose oxidation and NADPH generation. BioRxiv.

39. Nikel PI, Chavarría M, Fuhrer T, Sauer U, de Lorenzo V. 2015. *Pseudomonas putida* KT2440 Strain Metabolizes Glucose through a Cycle Formed by Enzymes of the Entner-Doudoroff, Embden-Meyerhof-Parnas, and Pentose Phosphate Pathways. J Biol Chem 290:25920–25932.

40. Wetmore KM, Price MN, Waters RJ, Lamson JS, He J, Hoover CA, Blow MJ, Bristow J, Butland G, Arkin AP, Deutschbauer A. 2015. Rapid quantification of mutant fitness in diverse bacteria by sequencing randomly bar-coded transposons. MBio 6:e00306–15.

41. Price MN, Wetmore KM, Waters RJ, Callaghan M, Ray J, Liu H, Kuehl JV, Melnyk RA, Lamson JS, Suh Y, Carlson HK, Esquivel Z, Sadeeshkumar H, Chakraborty R, Zane GM, Rubin BE, Wall JD, Visel A, Bristow J, Blow MJ, Deutschbauer AM. 2018. Mutant phenotypes for thousands of bacterial genes of unknown function. Nature 557:503–509.

42. Guzik MW, Narancic T, Ilic-Tomic T, Vojnovic S, Kenny ST, Casey WT, Duane GF, Casey E, Woods T, Babu RP, Nikodinovic-Runic J, O’Connor KE. 2014. Identification and characterization of an acyl-CoA dehydrogenase from Pseudomonas putida KT2440 that shows preference towards medium to long chain length fatty acids. Microbiology (Reading, Engl) 160:1760–1771.

43. Hume AR, Nikodinovic-Runic J, O’Connor KE. 2009. FadD from Pseudomonas putida CA-3 is a true long-chain fatty acyl coenzyme A synthetase that activates phenylalkanoic and alkanoic acids. J Bacteriol 191:7554–7565.

44. McMahon B, Mayhew SG. 2007. Identification and properties of an inducible phenylacyl-CoA dehydrogenase in Pseudomonas putida KT2440. FEMS Microbiol Lett 273:50–57.

45. Leščić Ašler I, Ivić N, Kovačić F, Schell S, Knorr J, Krauss U, Wilhelm S, Kojić-Prodić B, Jaeger K-E. 2010. Probing enzyme promiscuity of SGNH hydrolases. Chembiochem 11:2158–2167.

46. McMahon B, Gallagher ME, Mayhew SG. 2005. The protein coded by the PP2216 gene of Pseudomonas putida KT2440 is an acyl-CoA dehydrogenase that oxidises only short-chain aliphatic substrates. FEMS Microbiol Lett 250:121–127.

47. del Peso-Santos T, Bartolomé-Martín D, Fernández C, Alonso S, García JL, Díaz E, Shingler V, Perera J. 2006. Coregulation by phenylacetyl-coenzyme A-responsive PaaX integrates control of the upper and lower pathways for catabolism of styrene by Pseudomonas sp. strain Y2. J Bacteriol 188:4812–4821.

48. Ferrández A, García JL, Díaz E. 2000. Transcriptional regulation of the divergent paa catabolic operons for phenylacetic acid degradation in Escherichia coli. J Biol Chem 275:12214–12222.

49. Görisch H. 2003. The ethanol oxidation system and its regulation in Pseudomonas aeruginosa. Biochimica et Biophysica Acta (BBA) - Proteins and Proteomics 1647:98–102.

50. Hempel N, Görisch H, Mern DS. 2013. Gene ercA, encoding a putative iron-containing alcohol dehydrogenase, is involved in regulation of ethanol utilization in Pseudomonas aeruginosa. J Bacteriol 195:3925–3932.

51. Wehrmann M, Berthelot C, Billard P, Klebensberger J. 2018. The PedS2/PedR2 Two-Component System Is Crucial for the Rare Earth Element Switch in Pseudomonas putida KT2440. mSphere 3.

52. Arias S, Olivera ER, Arcos M, Naharro G, Luengo JM. 2008. Genetic analyses and molecular characterization of the pathways involved in the conversion of 2-phenylethylamine and 2-phenylethanol into phenylacetic acid in Pseudomonas putida U. Environ Microbiol 10:413–432.

53. Fernández M, Conde S, de la Torre J, Molina-Santiago C, Ramos J-L, Duque E. 2012. Mechanisms of resistance to chloramphenicol in Pseudomonas putida KT2440. Antimicrob Agents Chemother 56:1001–1009.

54. Gliese N, Khodaverdi V, Görisch H. 2010. The PQQ biosynthetic operons and their transcriptional regulation in Pseudomonas aeruginosa. Arch Microbiol 192:1–14.

55. García-Hidalgo J, Brink DP, Ravi K, Paul CJ, Lidén G, Gorwa-Grauslund MF. 2020. Vanillin Production in Pseudomonas: Whole-Genome Sequencing of Pseudomonas sp. Strain 9.1 and Reannotation of Pseudomonas putida CalA as a Vanillin Reductase. Appl Environ Microbiol 86.

56. Heeb S, Haas D. 2001. Regulatory roles of the GacS/GacA two-component system in plant-associated and other gram-negative bacteria. Mol Plant Microbe Interact 14:1351–1363.

57. Venturi V. 2003. Control of rpoS transcription in Escherichia coli and Pseudomonas: why so different? Mol Microbiol 49:1–9.

58. Bentley GJ, Narayanan N, Jha RK, Salvachúa D, Elmore JR, Peabody GL, Black BA, Ramirez K, De Capite A, Michener WE, Werner AZ, Klingeman DM, Schindel HS, Nelson R, Foust L, Guss AM, Dale T, Johnson CW, Beckham GT. 2020. Engineering glucose metabolism for enhanced muconic acid production in Pseudomonas putida KT2440. Metab Eng 59:64–75.

59. Ryan WJ, O’Leary ND, O’Mahony M, Dobson ADW. 2013. GacS-dependent regulation of polyhydroxyalkanoate synthesis in Pseudomonas putida CA-3. Appl Environ Microbiol 79:1795–1802.

60. Jacob K, Rasmussen A, Tyler P, Servos MM, Sylla M, Prado C, Daniele E, Sharp JS, Purdy AE. 2017. Regulation of acetyl-CoA synthetase transcription by the CrbS/R two-component system is conserved in genetically diverse environmental pathogens. PLoS ONE 12:e0177825.

61. Price MN, Arkin AP. 2017. PaperBLAST: Text Mining Papers for Information about Homologs. mSystems 2.

62. Monteagudo-Cascales E, García-Mauriño SM, Santero E, Canosa I. 2019. Unraveling the role of the CbrA histidine kinase in the signal transduction of the CbrAB two-component system in Pseudomonas putida. Sci Rep 9:9110.

63. Valentini M, García-Mauriño SM, Pérez-Martínez I, Santero E, Canosa I, Lapouge K. 2014. Hierarchical management of carbon sources is regulated similarly by the CbrA/B systems in Pseudomonas aeruginosa and Pseudomonas putida. Microbiology (Reading, Engl) 160:2243–2252.

64. Werle P, Morawietz M, Lundmark S, Sörensen K, Karvinen E, Lehtonen J. 2000. Alcohols, Polyhydric, p. *In* Wiley-VCH Verlag GmbH & Co. KGaA (ed.), Ullmann’s encyclopedia of industrial chemistry. Wiley-VCH Verlag GmbH & Co. KGaA, Weinheim, Germany.

65. Wang Y, Lv M, Zhang Y, Xiao X, Jiang T, Zhang W, Hu C, Gao C, Ma C, Xu P. 2014. Reconstruction of lactate utilization system in Pseudomonas putida KT2440: a novel biocatalyst for l-2-hydroxy-carboxylate production. Sci Rep 4:6939.

66. Jiang T, Guo X, Yan J, Zhang Y, Wang Y, Zhang M, Sheng B, Ma C, Xu P, Gao C. 2017. A Bacterial Multidomain NAD-Independent d-Lactate Dehydrogenase Utilizes Flavin Adenine Dinucleotide and Fe-S Clusters as Cofactors and Quinone as an Electron Acceptor for d-Lactate Oxidization. J Bacteriol 199.

67. Feller C, Günther R, Hofmann H-J, Grunow M. 2006. Molecular basis of substrate recognition in D-3-hydroxybutyrate dehydrogenase from Pseudomonas putida. Chembiochem 7:1410–1418.

68. Atsumi S, Hanai T, Liao JC. 2008. Non-fermentative pathways for synthesis of branched-chain higher alcohols as biofuels. Nature 451:86–89.

69. Conrad RS, Massey LK, Sokatch JR. 1974. D- and L-isoleucine metabolism and regulation of their pathways in Pseudomonas putida. J Bacteriol 118:103–111.

70. Roberts CM, Conrad RS, Sokatch JR. 1978. The role of enoyl-coa hydratase in the metabolism of isoleucine by Pseudomonas putida. Arch Microbiol 117:99–108.

71. Kang A, George KW, Wang G, Baidoo E, Keasling JD, Lee TS. 2016. Isopentenyl diphosphate (IPP)-bypass mevalonate pathways for isopentenol production. Metab Eng 34:25–35.

72. Sasaki Y, Eng T, Herbert RA, Trinh J, Chen Y, Rodriguez A, Gladden J, Simmons BA, Petzold CJ, Mukhopadhyay A. 2019. Engineering Corynebacterium glutamicum to produce the biogasoline isopentenol from plant biomass hydrolysates. Biotechnol Biofuels 12:41.

73. Hammer SK, Zhang Y, Avalos JL. 2020. Mitochondrial Compartmentalization Confers Specificity to the 2-Ketoacid Recursive Pathway: Increasing Isopentanol Production in Saccharomyces cerevisiae. ACS Synth Biol 9:546–555.

74. LaBauve AE, Wargo MJ. 2012. Growth and laboratory maintenance of Pseudomonas aeruginosa. Curr Protoc Microbiol Chapter 6:Unit 6E.1.

75. Ham TS, Dmytriv Z, Plahar H, Chen J, Hillson NJ, Keasling JD. 2012. Design, implementation and practice of JBEI-ICE: an open source biological part registry platform and tools. Nucleic Acids Res 40:e141.

76. Chen J, Densmore D, Ham TS, Keasling JD, Hillson NJ. 2012. DeviceEditor visual biological CAD canvas. J Biol Eng 6:1.

77. Hillson NJ, Rosengarten RD, Keasling JD. 2012. j5 DNA assembly design automation software. ACS Synth Biol 1:14–21.

78. Gibson DG, Young L, Chuang R-Y, Venter JC, Hutchison CA, Smith HO. 2009. Enzymatic assembly of DNA molecules up to several hundred kilobases. Nat Methods 6:343–345.

79. Engler C, Kandzia R, Marillonnet S. 2008. A one pot, one step, precision cloning method with high throughput capability. PLoS ONE 3:e3647.

80. Shanks RMQ, Kadouri DE, MacEachran DP, O’Toole GA. 2009. New yeast recombineering tools for bacteria. Plasmid 62:88–97.

81. Thompson MG, Pearson AN, Barajas JF, Cruz-Morales P, Sedaghatian N, Costello Z, Garber ME, Incha MR, Valencia LE, Baidoo EEK, Martin HG, Mukhopadhyay A, Keasling JD. 2020. Identification, Characterization, and Application of a Highly Sensitive Lactam Biosensor from Pseudomonas putida. ACS Synth Biol 9:53–62.

82. George KW, Thompson MG, Kang A, Baidoo E, Wang G, Chan LJG, Adams PD, Petzold CJ, Keasling JD, Lee TS. 2015. Metabolic engineering for the high-yield production of isoprenoid-based C₅ alcohols in E. coli. Sci Rep 5:11128.

83. Jones E, Oliphant T, Peterson P, Others. SciPy: Open source scientific tools for Python.

84. van der Walt S, Colbert SC, Varoquaux G. 2011. The NumPy Array: A Structure for Efficient Numerical Computation. Comput Sci Eng 13:22–30.

85. Kalyaanamoorthy S, Minh BQ, Wong TKF, von Haeseler A, Jermiin LS. 2017. ModelFinder: fast model selection for accurate phylogenetic estimates. Nat Methods 14:587–589.

86. Cruz-Morales P, Ramos-Aboites HE, Licona-Cassani C, Selem-Mójica N, Mejía-Ponce PM, Souza-Saldívar V, Barona-Gómez F. 2017. Actinobacteria phylogenomics, selective isolation from an iron oligotrophic environment and siderophore functional characterization, unveil new desferrioxamine traits. FEMS Microbiol Ecol 93.

